# CaMK1D signaling in AgRP neurons promotes ghrelin-mediated food intake

**DOI:** 10.1101/2021.12.07.471546

**Authors:** Kevin Vivot, Gergö Meszaros, Zhirong Zhang, Eric Erbs, Gagik Yeghiazaryan, Mar Quiñones, Erwan Grandgirard, Anna Schneider, Etienne Clauss--Creusot, Alexandre Charlet, Maya Faour, Claire Martin, Serge Luquet, Peter Kloppenburg, Ruben Nogueiras, Romeo Ricci

## Abstract

Hypothalamic AgRP/NPY neurons are key players in the control of feeding behavior. Ghrelin, a major hormone released under fasting conditions, activates orexigenic AgRP/NPY neurons to stimulate food intake and adiposity. However, cell-autonomous ghrelin-dependent signaling mechanisms in AgRP/NPY neurons remain poorly defined. Here we demonstrate that calcium/calmodulin-dependent protein kinase ID (CaMK1D), a genetic hot spot in type 2 diabetes, is activated in hypothalamus upon ghrelin stimulation and acts in AgRP neurons to promote ghrelin-dependent food intake. Global *CaMK1D* knockout mice are resistant to the orexigenic action of ghrelin, gain less body weight and are protected against high-fat diet-induced obesity. Deletion of *CaMK1D* in AgRP but not in POMC neurons is sufficient to recapitulate above phenotypes. Lack of CaMK1D attenuates phosphorylation of CREB and CREB-dependent expression of the orexigenic neuropeptides AgRP/NPY as well as the amount of AgRP fiber projections to the Paraventricular nucleus (PVN), while electrical activity of AgRP neurons and 5’ AMP-activated protein kinase (AMPK) signaling are unaffected. Hence, CaMK1D links ghrelin action to transcriptional control of orexigenic neuropeptide availability in AgRP neurons.

**Highlights:**

- Whole-body deletion of *CaMK1D* in mice reduces food intake, ghrelin sensitivity and protects against obesity.
- AgRP/NPY neuron-specific deletion of *CaMK1D* reduces food intake, ghrelin sensitivity, energy expenditure and protects against obesity.
- CaMK1D is dispensable for ghrelin-stimulated electrical activity of AgRP neurons and hypothalamic AMPK signaling.
- CaMK1D controls phosphorylation of CREB and CREB-dependent expression of the orexigenic neuropeptides AgRP and NPY.

**Graphical Abstract:** 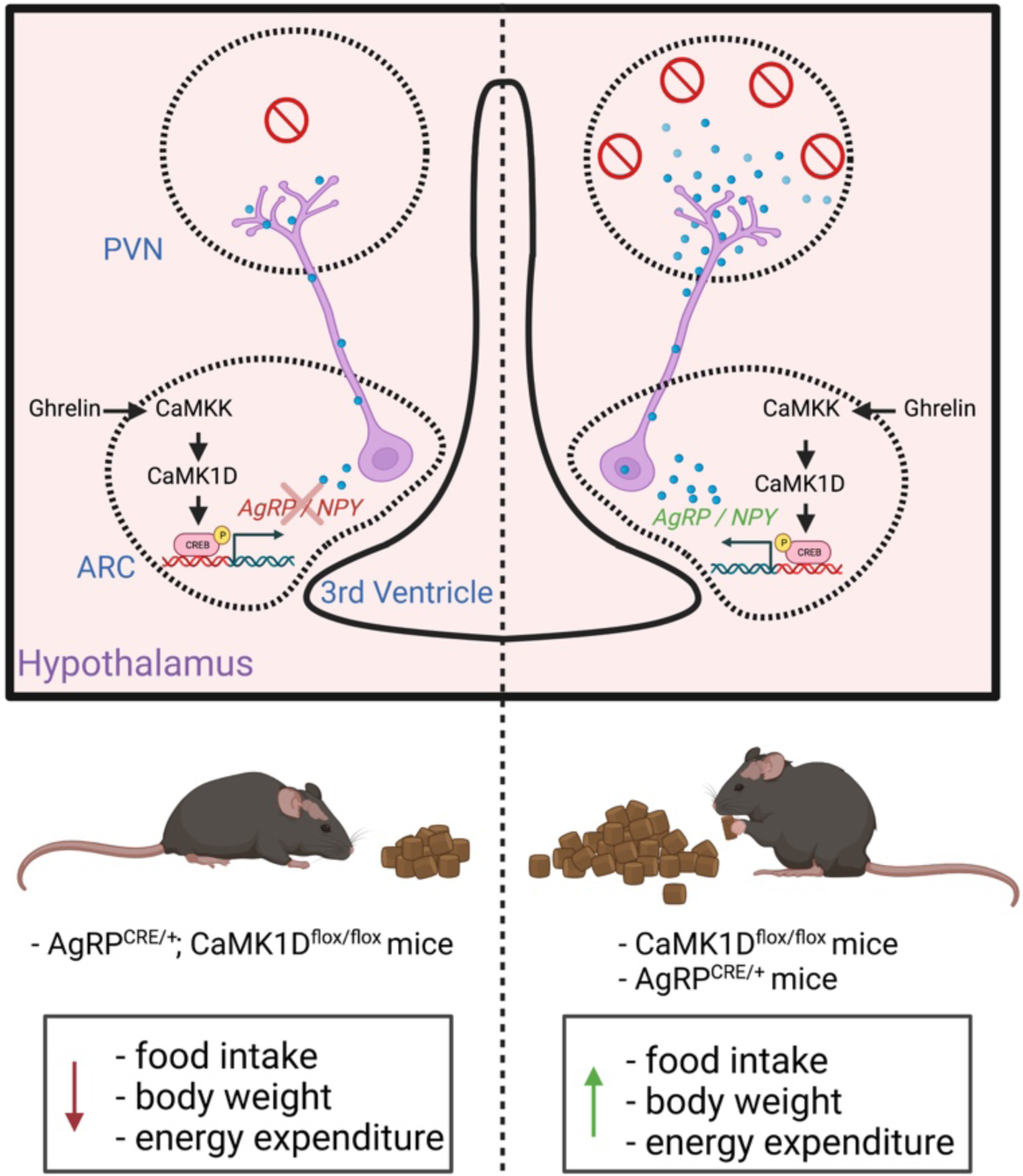

## Introduction

Tight regulation of energy homeostasis at multiple levels is instrumental for organisms to cope with changes in food availability. The Central Nervous System (CNS) orchestrates a complex array of processes mediating energy intake and expenditure. Hormonal, neuronal and nutritional signals according to changes in food absorption, energy storage and energy consumption in different organs reach the CNS which in turn triggers corresponding changes in feeding behavior and peripheral cellular metabolism (Kim et al., 2018).

Sensing of the nutrient status of the organism is governed by distinct neuronal cell populations, particularly within the arcuate nucleus (ARC) of the hypothalamus (Jais and Brüning, 2021). Neurons in this region provide specific projections to other hypothalamic nuclei including the paraventricular nucleus of the hypothalamus (PVN) or to different extrahypothalamic brain regions that in turn coordinate corresponding behavioral responses (Morton et al., 2006). Conversely, hypothalamic nuclei receive inputs from extrahypothalamic brain regions to control food intake and energy expenditure (Waterson and Horvath, 2015). Hypothalamic neurons also obtain signals from the mesolimbic reward system governing “hedonic” aspects of food intake (Stuber and Wise, 2016).

Orexigenic neuropeptide Y (NPY) and agouti-related peptide (AgRP)-expressing AgRP/NPY neurons and anorexigenic proopiomelancortin (POMC)-expressing neurons in the arcuate nucleus of the hypothalamus are primarily involved in the regulation of energy homeostasis. To control appetite and peripheral metabolism, these neurons are regulated by several hormones. Among others, leptin, ghrelin and insulin emerged as key players in this context. Both leptin and insulin receptors are expressed in these neurons and both insulin and leptin have been found to activate POMC and to inhibit AgRP/NPY neurons (Timper and Brüning, 2017). Ghrelin enhances the activity of AgRP/NPY neurons via its receptors, while it decreases the action of POMC neurons through a ghrelin receptor-independent mechanism (Chen et al., 2017).

Dysfunction of these neuronal circuits is known to contribute to overnutrition and obesity that eventually culminates in metabolic disorders such as type 2 diabetes (T2D) and/or cardiovascular diseases. Obesity and T2D are interlinked and complex metabolic disorders. Recent Genome-wide association studies (GWAS) and GWAS meta-analyses revealed complex polygenic factors influencing the development of both diseases. In fact more than ∼250 genetic loci have been identified for monogenic, syndromic, or common forms of T2D and/or obesity-related traits (Bonnefond and Froguel, 2015; Locke et al., 2015). Despite these remarkable advancements, the contribution of most obesity- and T2D-associated single nucleotide polymorphism (SNPs) to the pathogenesis of these diseases remains largely elusive.

CDC123 (cell division cycle protein 123)/CaMK1D (calcium/calmodulin-dependent protein kinase ID) represents one such locus on chromosome 10, strongly associated with T2D in European and Asian populations (Kooner et al., 2011; Shu et al., 2010; Zeggini et al., 2008). Among other variants, fine mapping identified rs11257655 as the predominant SNP within this locus (Morris et al., 2012). The change from C to T in the enhancer region of the rs11257655 allele, promotes DNA hypomethylation and the binding of the transcription factors FOXA1/FOXA2 on the enhancer region resulting in enhanced CaMK1D gene transcription (Thurner et al., 2018; Xue et al., 2018). Thus, CaMK1D expression might be enhanced and contributes to the development of T2D.

CaMK1D (or CaMK1δ) is the fourth member of the CaMK1 subfamily. CAMK1 has been described to be ubiquitously expressed at low levels in most tissues. Importantly however, CAMKI proteins are highly expressed in several brain regions, including the cortex, hippocampus, thalamus, hypothalamus, midbrain, and olfactory bulb, with each isoform exhibiting distinct spatiotemporal expression during neuronal development (Kamata et al., 2007). Ca^2+^/Calmodulin (CaM) binding and subsequent phosphorylation at Thr180 by CaMKK are required for CaMKI activation. In the particular case of CaMK1D, the phosphorylation of CaMK1D by CaMKK, induces a resistance to protein phosphatases, which keeps CaMK1D in a ‘primed’ state, to facilitate its activation in response to Ca^2+^ signals (Senga et al., 2015).

CaMKIs have been mainly shown to be important in neurons. CaMKIs control neuron morphology (Buchser et al., 2010) including axonal extension, growth cone motility (Wayman et al., 2004) and dendritogenesis (Takemoto-Kimura et al., 2007). It has been demonstrated that CaMKIs may also regulate neuronal function by controlling the long-term potentiation, a process involving persistent strengthening of synapses that leads to a long-lasting increase in signal transmission between neurons (Schmitt et al., 2005). However, specific non-redundant neuronal functions of any of the four members of the CaMKI subfamily including CaMK1D have yet to be determined. In fact, CaMK1D has been shown to be implicated in neutrophil function (Verploegen et al., 2005). The genetic association of the *CaMK1D* locus with T2D and correlation of clinical data revealed that CaMK1D might promote pancreatic β cell dysfunction (Simonis-Bik et al., 2010). However, direct experimental evidence for the latter conclusion is lacking thus far. Another study proposed that CaMK1D stimulates hepatic glucose output, a mechanism contributing to T2D (Haney et al., 2013).

Using global and conditional CaMK1D knockout mice, we provide evidence for a role of CaMK1D in central regulation of food intake, while its function seems to be redundant in the liver and in the pancreatic β cell. We demonstrate that CaMK1D acts in hypothalamic AgRP neurons to control food intake in response to ghrelin. While CaMK1D in AgRP neurons is redundant for ghrelin-stimulated increase in electrical neuronal activity and AMPK signaling, its absence impairs ghrelin-induced activatory CREB phosphorylation, AgRP transcription and AgRP/NPY fiber abundance within the PVN. Our data thus unveil ghrelin signaling mechanisms in AgRP neurons downstream or independent of neuronal activation to be necessary for efficient appetite stimulation.

## Results

### Global deletion of CaMK1D in mice protects against obesity

To understand the function of CaMK1D in metabolism, we generated mice carrying a floxed allele of *CaMK1D* (Figure S1A) and crossed them with the global Cre-deleter, Rosa26-Cre (Soriano, 1999), to obtain whole-body knockout mice. Western blotting confirmed efficient deletion of CaMK1D in different organs including brain, pancreas and intestine (Figure 1A). Whole-body *CaMK1D* knockout mice (*CaMK1D^-/-^* mice) were born in expected gender distribution (Figure S1B) and Mendelian ratios (Figure S1C) and developed without overt problems. Body and tibia lengths at 7 weeks of age were equal in *CaMK1D^-/-^* and *CaMK1D WT* mice (*CaMK1D^+/+^* mice) (Figure S1D and S1E), thus excluding any major postnatal growth defect. However, body weight gain in *CaMK1D^-/-^* mice on a chow diet was significantly reduced from 16 weeks on after starting to measure weight at the age of 5 weeks as compared to *CaMK1D^+/+^* mice. This significant difference was exacerbated when obesity was induced in mice on a high fat diet (HFD) at the age of 5 weeks (Figure 1B). Quantitative Nuclear Magnetic Resonance (qNMR) revealed that HFD-fed *CaMK1D^-/-^* mice compared to *CaMK1D^+/+^* mice had a reduced fat mass, while there was no significant difference in the lean mass and free body fluid (FBF) (Figure 1C). In line with reduced obesity, fasted glucose levels at 10 weeks after HFD feeding in *CaMK1D^-/-^* mice was significantly reduced as compared to *CaMK1D^+/+^* mice (Figure 1D). In fact, glucose levels in *CaMK1D^-/-^* mice on a HFD were similar as in corresponding chow diet-fed mice indicating that they were protected from obesity-induced hyperglycemia. There was no apparent difference in fasting glucose levels between *CaMK1D^-/-^* and *CaMK1D^+/+^* mice on a chow diet (Figure 1D). The observed reduced fasting glucose levels correlated with reduced fasting insulin levels (Figure 1E). Glucose tolerance was slightly, but not significantly, improved in *CaMK1D^-/-^* as compared to CaMK1D^+/+^ mice on a chow diet (Figure 1F). However, these differences became significant in mice on a HFD (Figure 1G and 1H). While insulin sensitivity was unaltered in *CaMK1D^-/-^* mice on a chow diet (Figure 1I), it was significantly improved in *CaMK1D^-/-^* as compared to *CaMK1D^+/+^* mice on a HFD (Figure 1J and 1K). Glucose-induced secretion of insulin (GSIS) was slightly but not significantly reduced in *CaMK1D^-/-^* mice as compared to *CaMK1D^+/+^* mice on a HFD (Figure S2A). GSIS from isolated islets of *CaMK1D^-/-^* and *CaMK1D^+/+^* mice was comparable (Figure S2B).

**Figure 1.**
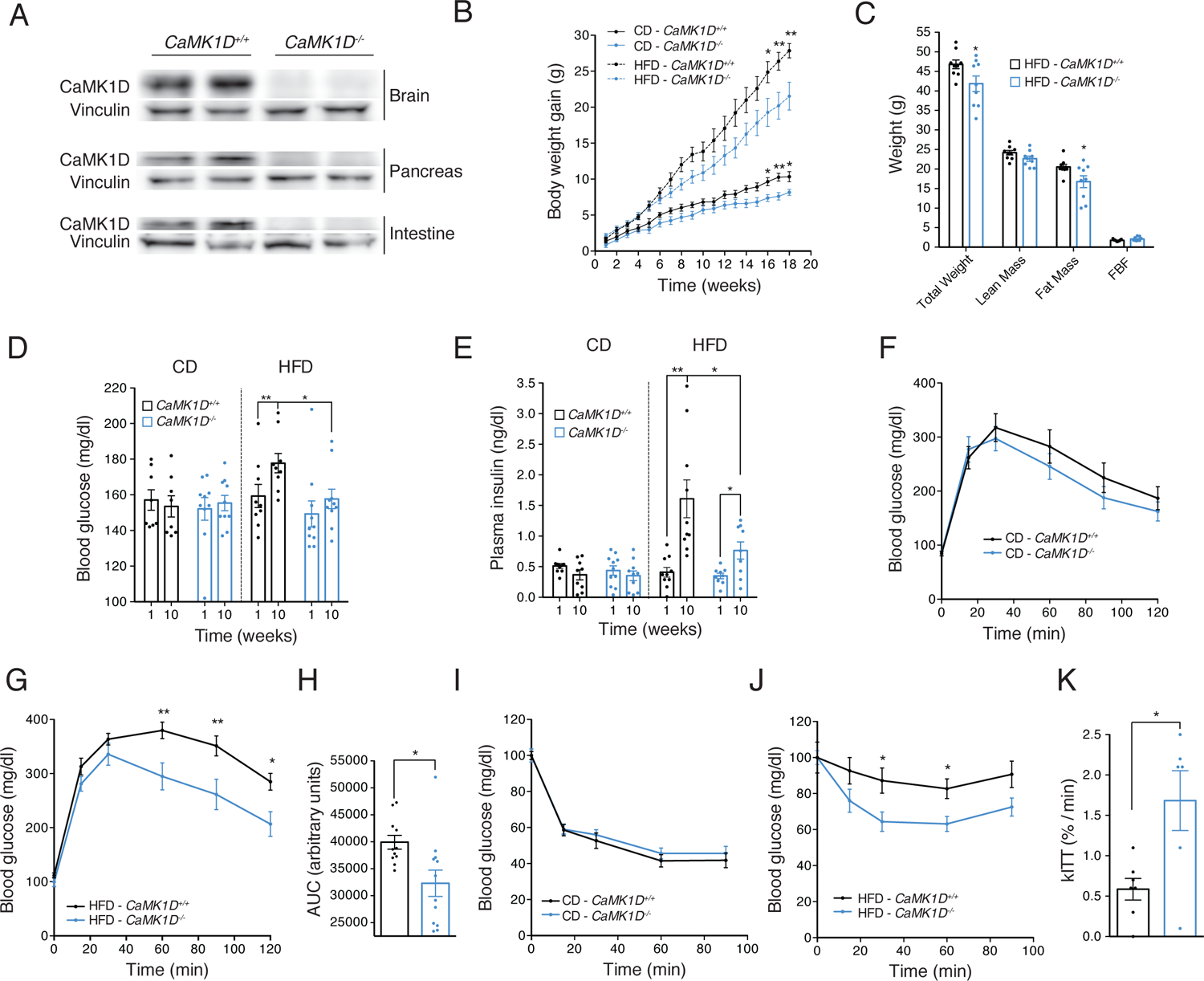
Global Deletion of *CaMK1D* gene reduces obesity in mice. (A) Expression of CaMK1D protein in different tissues as indicated. Tissues from the wild type (*CaMK1D^+/+^*) and whole-body *CaMK1D* knockout (*CaMK1D^-/-^*) mice were analyzed for CaMK1D expression by western blot. (B) Body weight gain of mice with indicated genotypes fed with a Chow diet (CD) or a High Fat Diet (HFD). (n=9 to 11/ group). (C) Body composition of mice with indicated genotypes fed with a HFD measured by qNMR. (n=9/ group) (*p ≤ 0.05) (FBF = free body fluid). (D) 4 hours fasted blood glucose levels at indicated time points. (n=8 to 10 / group). (E) 4 hours fasted plasma insulin levels at indicated time points. (n=9 to 11 / group). (F-H) Blood glucose levels during an IPGTT in mice with the indicated genotypes and diets. IPGTT was performed after an overnight food withdrawal. Corresponding areas Under the Curve (AUC) are depicted. (n=9 to 12 / group). (I-K) Blood glucose levels during ITT in mice with the indicated genotypes and diets. IPGTTs were performed after a 4H food withdrawal. kITTs obtained from ITT are depicted. (n=6 to 10 / group). ∗p < 0.05 and ∗∗p < 0.01. Statistical tests included two-way ANOVA (B, H and J) and unpaired Student’s t test (C, D, E, H and K).

To further exclude a primary role of CaMK1D in the pancreas including β cells, we crossed floxed mice with *PDX1^Cre/+^* mice (Gu et al., 2002). Western blotting confirmed that deletion of *CaMK1D* in the pancreas was efficient, while no apparent deletion was visible in brain (Figure S2C) or other organs (data not shown). Pancreas-specific *CaMK1D* knockout mice (*PDX1^Cre/+^;CaMK1D^flox/flox^* mice) did not show any differences in body weight gain and glucose tolerance as compared to floxed control mice (*CaMK1D^flox/flox^* mice) on a chow diet as well as HFD (Figure S2D-F), confirming that CaMK1D is redundant in β cell function.

Hepatic insulin resistance leading to increased gluconeogenesis is an important mechanism contributing to obesity-related changes in glucose homeostasis. Therefore, we generated liver-specific knockout mice using *Albumin^Cre/+^* mice (Postic et al., 1999) (*Alb^Cre/+^;CaMK1D^flox/flox^* mice) and corresponding floxed control mice (*CaMK1D^flox/flox^* mice). Quantitative reverse transcription PCR (qRT-PCR) confirmed efficient deletion of CaMK1D in the liver (Figure S2G) but not in other organs (data not shown). However, hepatic deletion of *CaMK1D* did not affect body weight gain, glucose tolerance and insulin sensitivity in mice on a chow diet as well as on HFD (Figure S2H-L). Thus, our data exclude any major functions of CaMK1D in pancreatic β cells and liver under both normal and obesity conditions.

### Global deletion of CaMK1D alters ghrelin-mediated food intake

To further understand reduced body weight gain and fat mass in *CaMK1D^-/-^* mice as compared to *CaMK1D^+/+^* mice, we next explored energy metabolism. In line with reduced obesity, cumulative food intake was decreased in *CaMK1D^-/-^* mice as compared to *CaMK1D^+/+^* mice on a chow diet as well as on HFD reaching significance at 16 weeks after starting to measure it (Figure 2A). Likewise, cumulative food intake in response to 24 hours fasting was significantly reduced in *CaMK1D^-/-^* mice throughout the observed period of refeeding as compared to *CaMK1D^+/+^* mice (Figure 2B). Indirect calorimetry revealed that energy expenditure (Figure 2C and 2D), O_2_ consumption (Figure 2E) CO_2_ production (Figure 2F) and the respiration exchange rate (RER) (Figure 2G) were equal in *CaMK1D^-/-^* mice as compared to *CaMK1D^+/+^* mice on a HFD. Moreover, the locomotor activity under basal conditions was comparable in *CaMK1D^-/-^* mice and *CaMK1D^+/+^* mice (data not shown). However, locomotor activity of *CaMK1D^-/-^* as compared to *CaMK1D^+/+^* mice was significantly reduced in the night period after 24 hours fasting (Figure 2H and 2I) in line with reduced appetite and reduced seeking for food in response to fasting. Thus, reduced obesity primarily correlated with reduced appetite and food intake.

**Figure 2.**
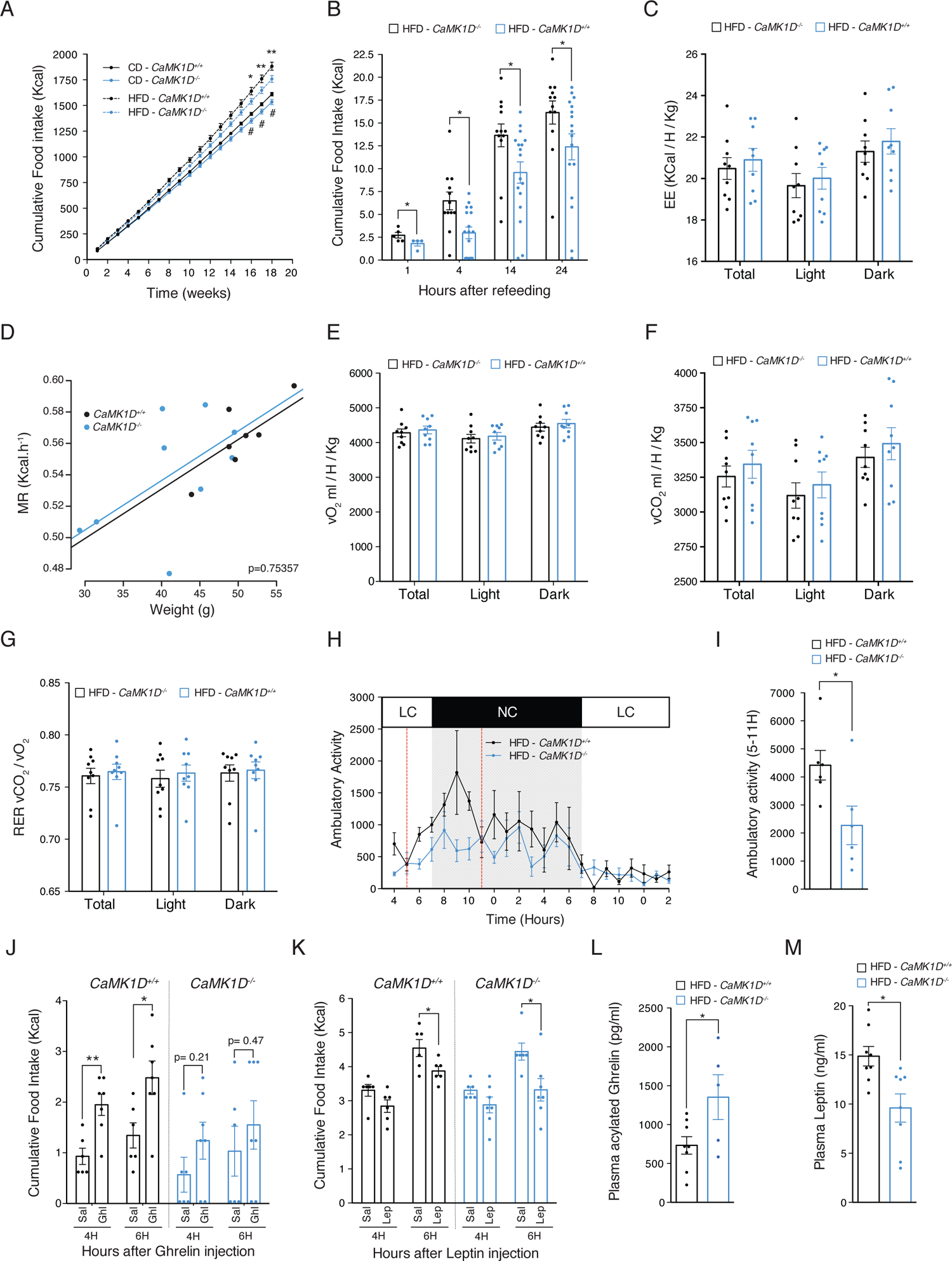
Deletion of *CaMK1D* attenuates ghrelin-induced food intake. (A) Cumulative food intake from mice with indicated genotypes and diets. (n=9 to 11 / group). (B) Cumulative food intake from mice with indicated genotypes on a HFD. Food intake was determined 24H after food withdrawal. (n=12 to 17 / group). (C) Energy expenditure, (D) Regression-based analysis of absolute MR against body mass. The ANCOVA analysis was done with MR as a dependent variable, the genotype as a fixed variable and body mass as a covariate. (E) consumed O_2_, (F) produced CO_2_ and (G) the respiratory exchange ratio (RER) in HFD fed control (*CaMK1D^+/+^)* and whole body *CaMK1D* knockout (*CaMK1D^-/-^*) mice (n=9 / group). (H-I) Ambulatory activity over 24H. Ambulatory activity was measured with mice deprived from food for 24H. AUC from 5PM to 11PM was calculated (n=6 / group). (J) Cumulative food intake after ghrelin injections (30 µg / day) of mice with indicated genotypes on a chow diet. (n=6 to 7 / group). (K) Cumulative food intake after leptin injections (3mg / kg) of mice with indicated genotypes on a chow diet. Food intake was determined 24H after food withdrawal. (n=6 to 7 / group). (L) Blood acylated Ghrelin and (M) Blood Leptin levels in mice with indicated genotypes on a HFD. Blood sampling was performed 4H after food withdrawal. (N=5 to 8 / group). ∗p < 0.05 and ∗∗p < 0.01. Statistical tests included two-way ANOVA (A, B), ANCOVA (D) and unpaired Student’s t test (I,J,K,L,M).

Ghrelin is a gut-derived hormone released in response to fasting and promotes feeding behavior and adiposity (Müller et al., 2015). Given that the resistance to diet-induced obesity of *CaMK1D^-/-^* mice could be explained by reduced food intake, we next wondered whether the ghrelin response was affected in mice lacking *CaMK1D*. To this end, we determined cumulative food intake upon intraperitoneal injections of ghrelin in mice on a chow diet. While *CaMK1D^+/+^* mice showed a significant increase in cumulative food intake at 4 and 6 hours after ghrelin injections, such a response was almost absent in *CaMK1D^-/-^* mice (Figure 2J). In contrast, the response to leptin was comparable in *CaMK1D^+/+^* mice and *CaMK1D^-/-^* mice (Figure 2K). Blood levels of acylated ghrelin were significantly higher in *CaMK1D^-/-^* as compared to *CaMK1D^+/+^* mice on a HFD (Figure 2L) in line with an adaptive response to a primary defect in ghrelin action. Conversely, blood levels of leptin were significantly lower in *CaMK1D^-/-^* mice as compared to *CaMK1D^+/+^* mice (Figure 2M) on a HFD, correlating well with the degree of obesity.

To exclude any major anxiety-like behavior or stress-induced anhedonia, we subjected mice to an open field and to a sucrose preference test, respectively. No major differences between genotypes could be observed (Figure S3A-H). Thus, lack of *CaMK1D* results in a compromised ghrelin response, which is in line with reduced obesity.

### CaMK1D acts in AgRP neurons to regulate food intake in response to ghrelin

Ghrelin stimulates central neurons to promote feeding. We thus next asked whether CaMK1D in the nervous system was mainly responsible for the observed phenotype in *CaMK1D^-/-^* mice. We therefore crossed *CaMK1D^flox/flox^* mice with *Nestin^Cre/+^* mice resulting in efficient deletion of *CaMK1D* in brain including hypothalamus but not in other organs such as intestine and pancreas (Figure S4A). Indeed, the body weight gain was significantly attenuated in nervous system-specific *CaMK1D* knockout mice (*Nestin^Cre/+^*;*CaMK1D^flox/flox^* mice) as compared to Cre and floxed control mice (*Nestin^Cre/+^* mice and *CaMK1D*^flox/flox^ mice) on a chow diet as well as on HFD (Figure S4B). Cumulative food intake was decreased in *Nestin^Cre/+^*;*CaMK1D^flox/flox^* mice as compared to control mice on a chow diet (Figure S4C) as well as on a HFD (Figure S4D). Likewise, cumulative food intake in response to 24 hours fasting was significantly reduced in *Nestin^Cre/+^*;*CaMK1D^flox/flox^* mice as compared to control mice after 4, 14 and 24 hours of refeeding (Figure S4E). Consistently, while control mice showed a significant increase in cumulative food intake at 4 and 6 hours after ghrelin injections, such a response was absent in *Nestin^Cre/+^*;*CaMK1D^flox/flox^* mice (Figure S4F).

Ghrelin primarily acts on hypothalamic neurons in the arcuate nucleus. In particular, it stimulates NPY/AgRP neurons to promote appetite (Andrews et al., 2008). Given our results in nervous system-specific knockout mice, we next asked whether CaMK1D acts in AgRP neurons to control food intake. To this end, we crossed *CaMK1D^flox/flox^* mice with *AgRP^Cre/+^* mice resulting in efficient recombination of the *CaMK1D* locus in hypothalamus but not in the brain cortex, liver, tail and white blood cells (Figure 3A). Strikingly, AgRP neuron-specific knockout mice (*AgRP^Cre/+^*; *CaMK1D^flox/flox^* mice) gained significantly less weight as compared to Cre and floxed control mice (*AgRP^Cre/+^* mice and *CaMK1D^flox/flox^* mice) on a chow diet as well as on HFD (Figure 3B). Similar to nervous system-specific knockout mice, *AgRP^Cre/+^*; *CaMK1D^flox/flox^* mice showed significantly less cumulative food intake as compared to control mice on a chow diet (Figure 3C) as well as HFD (Figure 3D). Similar significant differences were observed in the cumulative food intake in response to 24 hours fasting (Figure 3E) as well as in response to ghrelin injection (Figure 3F). Importantly, no such differences could be detected when deleting *CaMK1D* in anorexigenic POMC neurons (Figure S5A–F), suggesting that the effects of CaMK1D on food intake are specific to AgRP neurons.

**Figure 3.**
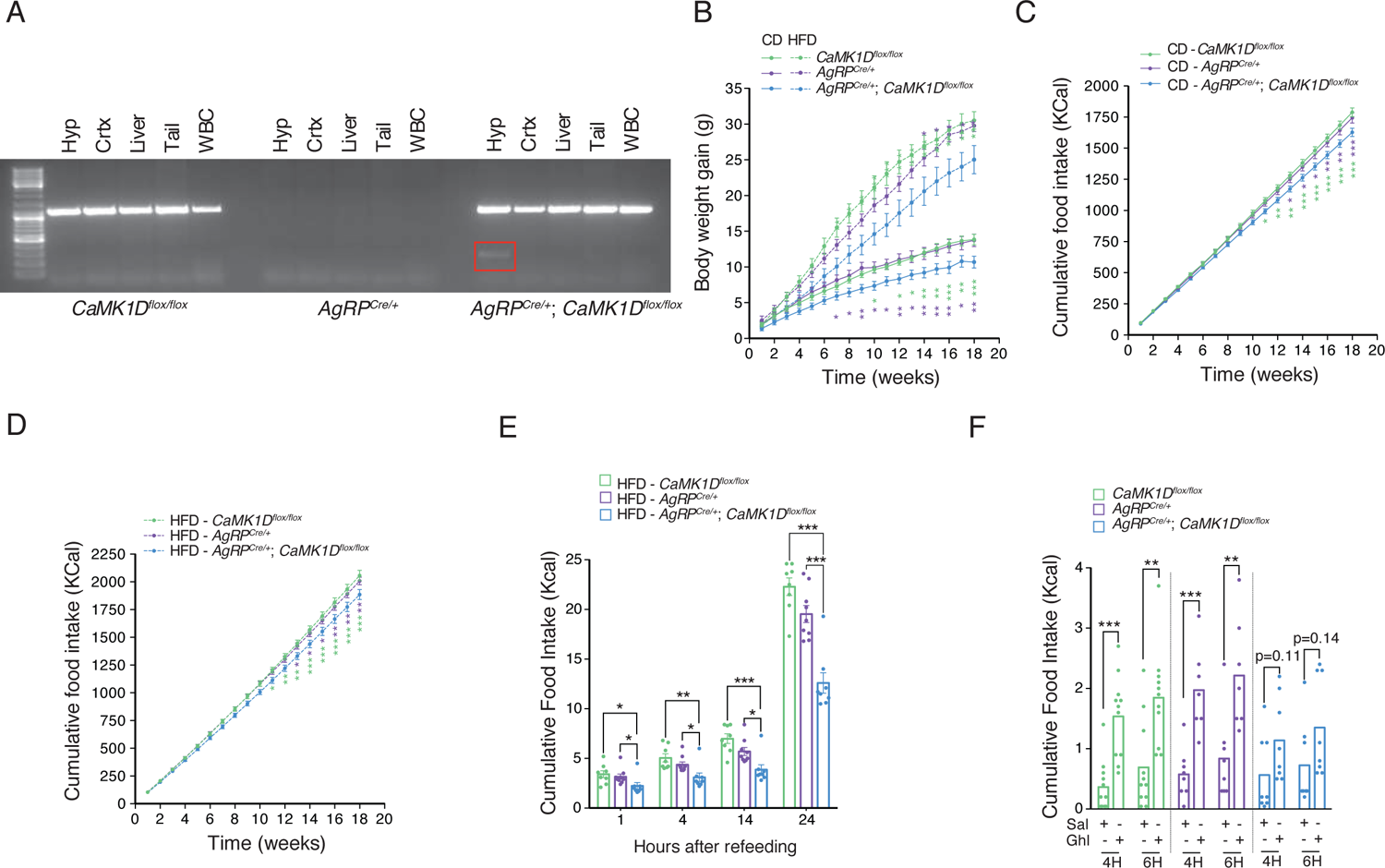
Conditional deletion of *CaMK1D* in AgRP neurons leads to reduced body weight gain and food intake. (A) Verification of recombination in the CaMK1D locus in hypothalamus (Hyp), cortex (Crtx), liver, tail and white blood cells (WBC) from mice with indicated genotypes. DNA from tissues were analyzed by PCR with primers amplifying either recombined or floxed alleles, respectively. (B) Body weight gain of mice with indicated genotypes on a Chow diet (CD) or on a High Fat Diet (HFD). (n=10 to 15/ group). (C-D) Cumulative food intake of mice with indicated genotypes fed on Chow diet or on high fat diet. (n=10 to 15 / group). (E) Cumulative food intake of mice with indicated genotypes and diets. Food intake was determined 24H after fasting. (n=12 to 17 / group). (F) Cumulative food intake after ghrelin injections of mice with indicated genotypes and diets. (n=6 to 7 / group). ∗p < 0.05, ∗∗p< 0.01 and ∗∗∗p < 0.001. Statistical tests included two-way ANOVA (B,C,D) and unpaired Student’s t test (E, F).

To further evaluate a role of CaMK1D in AgRP neuron-dependent energy metabolism, we next performed indirect calorimetry with *AgRP^Cre/+^*;*CaMK1D^flox/flox^* mice and control mice on a chow diet. Cumulative food intake (Figure 4A) and locomotor activity (Figure 4B and 4C) were comparable in *CaMK1D^-/-^* mice and *CaMK1D^+/+^* mice. Interestingly, energy expenditure (Figure 4D and 4E) was decreased in *AgRP^Cre/+^*;*CaMK1D^flox/flox^* mice as compared to control mice. The regression-based analysis-of-covariance (ANCOVA) confirmed that there was a body weight-independent metabolic rate (MR) difference with lower MR in the *AgRP^Cre/+^*;*CaMK1D^flox/flox^* relative to the control mice (Figure 4F). In line with these findings, O_2_ consumption (Figure 4G) and CO_2_ production (Figure 4H) were also reduced, while the respiration exchange rate (RER) (Figure 4I) was equal. Given the overall reduction in body weight gain, reduced energy expenditure was most likely compensatory to compromised energy availability caused by reduced food intake. Altogether, our data thus suggest that CaMK1D acts in AgRP neurons to primarily control food intake in response to ghrelin.

**Figure 4.**
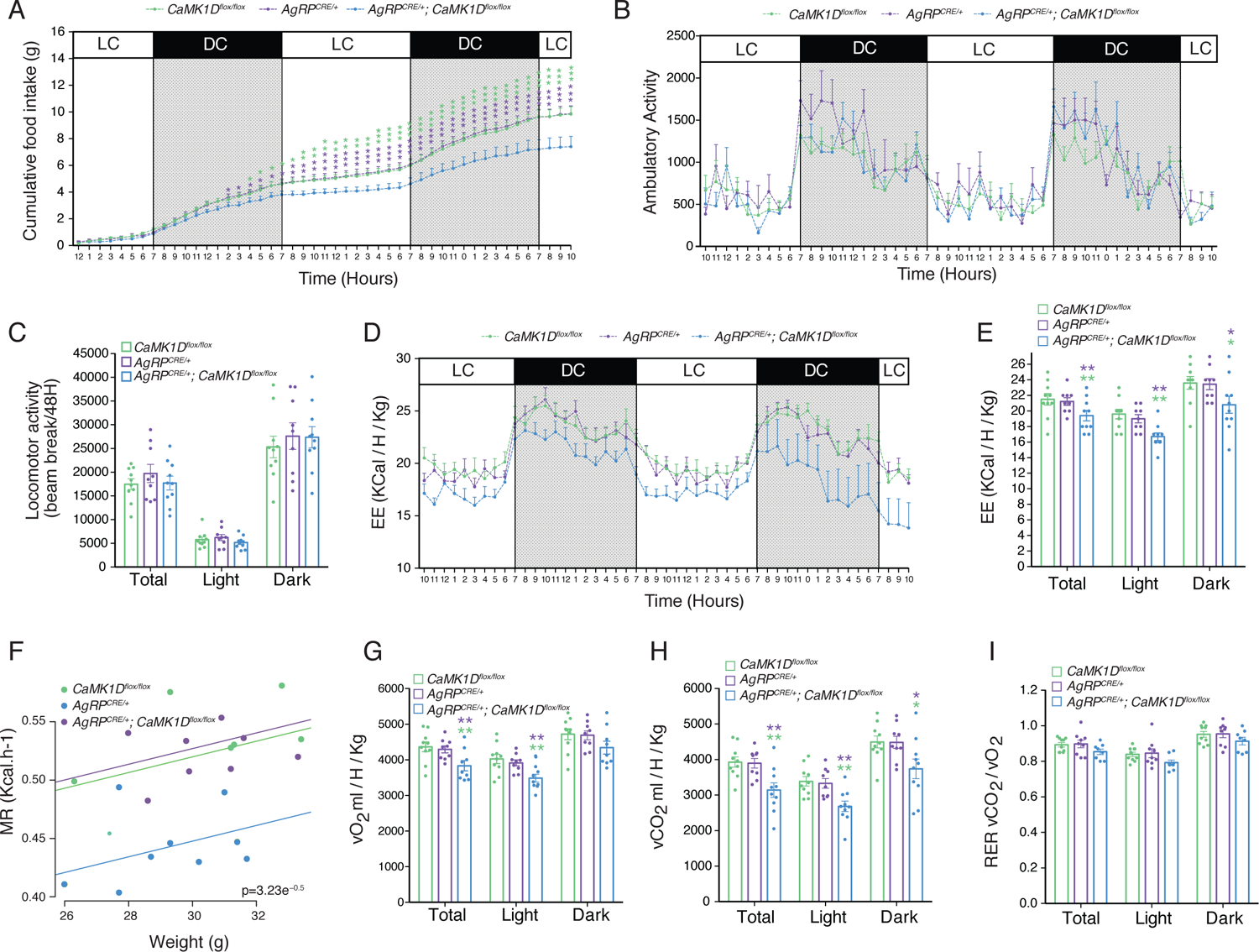
Conditional deletion of *CaMK1D* in AgRP neurons decreases energy expenditure. (A) Cumulative food intake from mice with indicated genotypes on a CD. Food intake was determined 48H during indirect calorimetric measurements. (N=10 to 11 / group). (B-C) Locomotor activity over 48H in mice with indicated genotypes on a Chow diet. (D-E) Energy expenditure, (F) Regression-based analysis of absolute MR against body mass. The ANCOVA analysis was done with MR as a dependent variable, the genotype as a fixed variable and body mass as a covariate. (G) consumed O_2_, (H) produced CO_2_ and (I) the respiratory exchange ratio (RER) in of mice with indicated genotypes on a Chow diet. All indirect calorimetric measurements were done in automated cages. (N=10-11/ group). ∗p < 0.05, ∗∗p< 0.01 and ∗∗∗p < 0.001. Statistical tests included two-way ANOVA (A), ANCOVA (F) and unpaired Student’s t test (E,G,H).

### Deletion of CaMK1D in mice does not affect AgRP/NPY neuronal activity in response to ghrelin

c-fos expression is used as a marker for neuronal activity (Hoffman et al., 1993). To understand the function of CaMK1D in ghrelin-induced neuronal activity, we next explored c-fos expression in AgRP neurons in the absence and presence of CaMK1D. Immunofluorescence of c-fos revealed no significant differences in basal and ghrelin-induced expression of c-fos in ARC neurons of *CaMK1D*^-/-^ mice as compared to control mice on a chow diet (Figure 5A and 5B). To verify this finding, we compared the effect of ghrelin on AgRP neurons between both mice lines using perforated patch clamp recordings in brain slices. To identify AgRP neurons, we generated *CaMK1D*^-/-^ and *CaMK1D^+/+^* mice carrying a EGFP reporter under the control of the *AgRP* promoter (*CaMK1D*^-/-^-AgRP-EGFP and *CaMK1D^+/+^*-AgRP-EGFP mice) (Figure 5C). The ghrelin-induced increases in action potential frequency were similar in AgRP neurons of *CaMK1D*^-/-^-AgRP-EGFP as compared to *CaMK1D^+/+^*-AgRP-EGFP mice (*CaMK1D^+/+^*, n=12; *CaMK1D*^-/-^, n=10, p=0.82) (Figure 5D and 5E). Between the two mouse lines, we also found no significant differences in general intrinsic electrophysiological properties of AgRP neurons such as spontaneous actions potential frequency, input resistance, excitability, and whole-cell capacitance (Figure 5F-I). Taken together, these data suggest that CaMK1D is redundant for ghrelin-stimulated activation of AgRP neurons.

**Figure 5.**
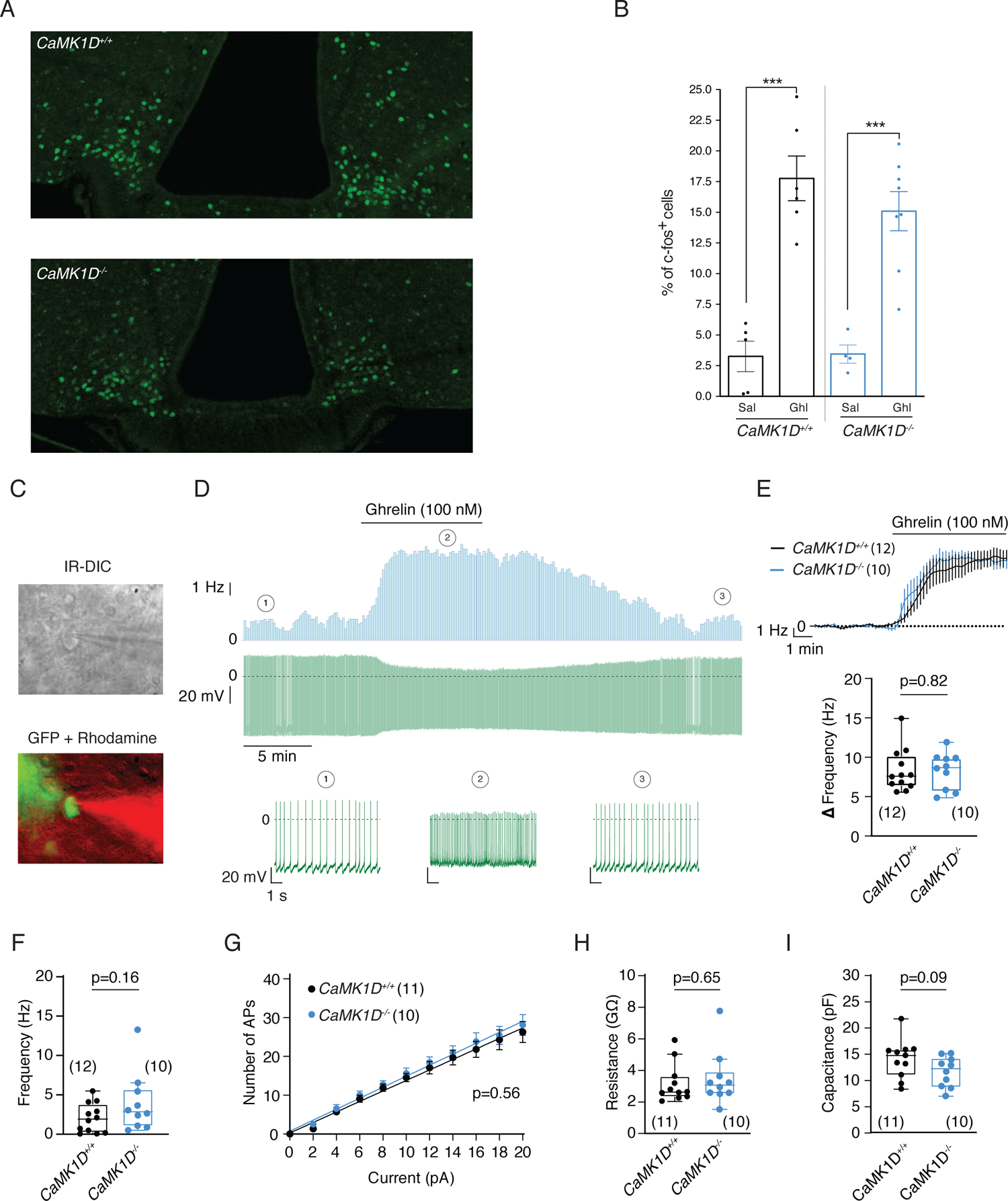
CaMK1D is dispensable for ghrelin-induced CaMK1D is dispensable for ghrelin-induced electrophysiological activation of AgRP/NPY neurons (A-B) Representative immunofluorescence and quantification of c-fos^+^ cells in the ARC of mice with indicated genotypes. Animals were injected with 30 µg ghrelin or vehicle and 2H after injections whole hypothalamus was removed. (C) Recording situation shown in a brightfield (top) and fluorescent image (bottom). The AgRP neuron expressed EGFP (green) and the recording pipette contained tetramethyl rhodamine dextrane (red) to monitor membrane integrity during the perforated patch clamp recording. (D) Recording of an AgRP neuron from a *CaMK1D^+/+^* mouse during ghrelin (100 nM) bath application. Top: Rate histogram, bin width: 10 s. Middle: Original recording. Bottom: Segments of the original recording in higher time resolution. The numbers indicate the time points from which the segments originate. (E) Ghrelin responses of AgRP neurons from *CaMK1D^+/+^* and in *CaMK1D^-/-^* mice, expressed as change in action potential frequency. Top: Mean (± SEM) responses during the first 5 min of ghrelin application. Bottom: Box plots showing the change in action potenial frequency measured between 6 and 8 min of ghrelin application. (F-I) Basic electrophysiological properties of AgRP neurons in *CaMK1D^+/+^* and in *CaMK1D^-/-^* mice. (F) Spontaneous action potential frequency. (G) Excitability assessed by the number of action potentials (APs) as a function of current pulse (1s) amplitude. (H) Input resistance. (I) Whole-cell capacitance. The horizontal lines in the box plots show the median. The whiskers were calculated according to the ‘Tukey’ method. ∗∗∗p < 0.001. Statistical tests included unpaired Student’s t test (A). Data in (E, F, H, and I) were compared using the Mann-Whitney-U-test, and linear regressions in (G) were compared using the F-test. n values are given in brackets.

### Ghrelin activates CaMK1D to induce AgRP/NPY expression and projections in the PVN

Lack of CaMK1D did not affect ghrelin-induced increase in electrical activity of AgRP neurons prompting us to hypothesize that CaMK1D acts downstream or independent of neuronal activity. We thus next asked whether CaMK1D is activated upon ghrelin. We first used Phos-tag gels to address CaMK1D phosphorylation in response to ghrelin in cultured primary hypothalamic cells isolated from *CaMK1D^-/-^* and *CaMK1D^+/+^* mice. Indeed, phosphorylated CaMK1D as marked by the upshifted detected band increased upon ghrelin stimulation (Figure 6A). Calcium/calmodulin directly activates calium/calmodulin-dependent protein kinase I by binding to the enzyme and indirectly promotes the phosphorylation and synergistic activation of the enzyme by calcium/calmodulin-dependent protein kinase kinase (CaMKK) (Haribabu et al., 1995). In line with above findings in cultured neurons, activatory phosphorylation of CaMK1D increased in hypothalamus of ghrelin-stimulated *CaMK1D^+/+^* mice, while no phosphorylation of CaMK1D or total protein was visible in *CaMK1D^-/-^* samples (Figure 6B).

**Figure 6.**
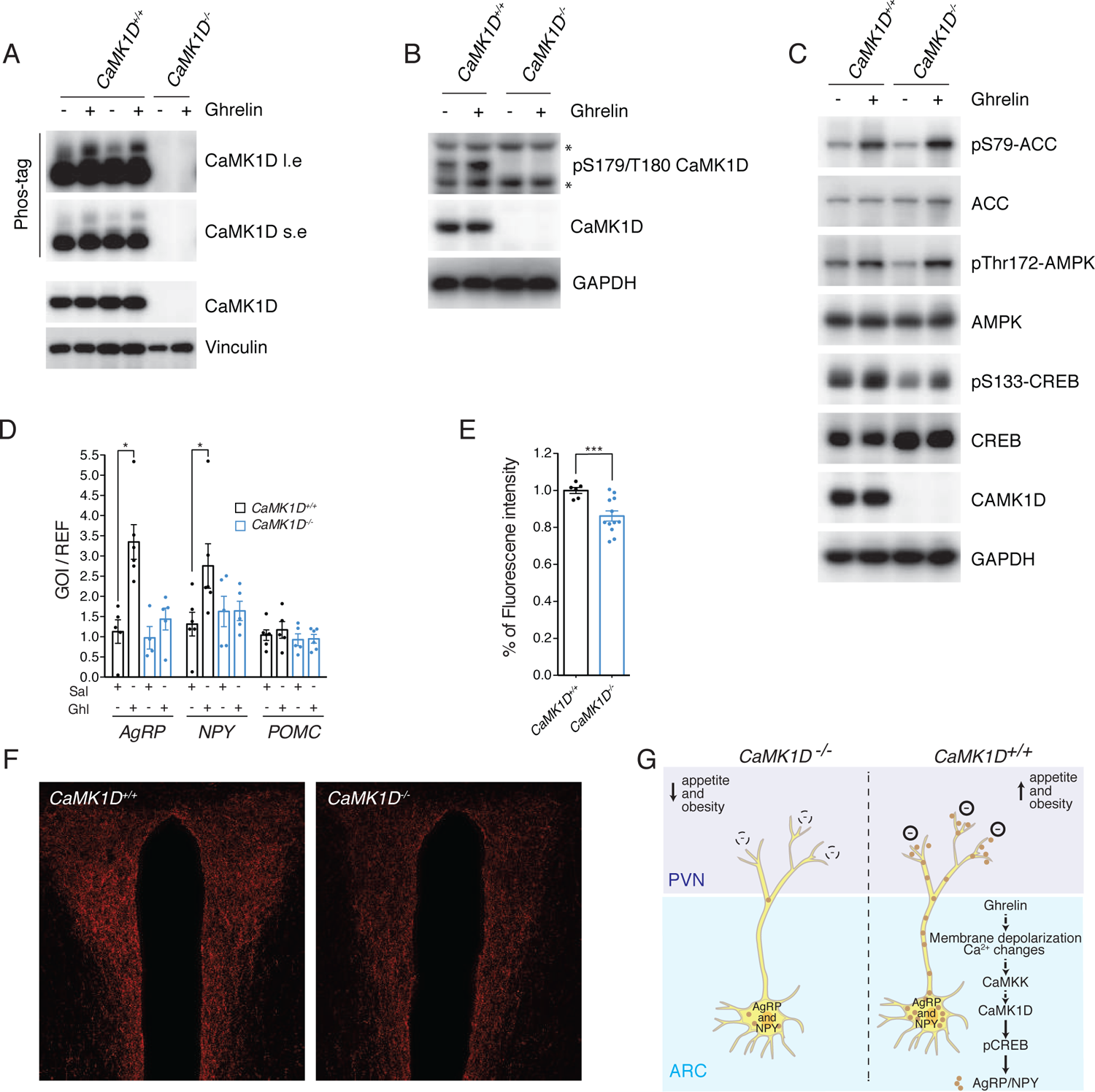
Lack of CaMK1D reduces ghrelin-induced AgRP/NPY expression and abundance in AgRP neuron projections to the PVN. (A) Representative Phos-tag immunoblot of CaMK1D using lysates of hypothalamic cells from mice with indicated genotypes treated with 1 µM Ghrelin or vehicle for 5 min. Vinculin was used as a loading control. (B) Representative immunoblot of pS179/T180 CaMK1D using lysates of whole hypothalamus from mice with indicated genotypes. Animals were injected with 30 µg ghrelin or vehicle and 2H after injections whole hypothalamus was removed and used for protein extraction. GAPDH was used as a loading control. (C) Representative immunoblots of pS79-ACC, pT172-AMPK and pS133-CREB using lysates of hypothalamic primary neurons from mice with indicated genotypes treated with 1 µM Ghrelin or vehicle for 5 min. GAPDH was used as a loading control. (D) Expression of AgRP, NPY and POMC mRNA in whole hypothalamus. Animals were injected with 30 µg ghrelin or vehicle and whole hypothalamus was removed 2H after injections. Tissues from mice with indicated genotypes were analyzed by qPCR (n=5 to 6 / group). (E-F) Representative immunofluorescence and quantification of AgRP projections to the PVN of mice with indicated genotypes. Animals were injected with 30 µg ghrelin and 2H after injections, whole hypothalamus was removed or brain was extracted and sliced. (G) Schematic model depicting mechanisms as two how CaMK1D promotes food intake. ∗p < 0.05 and ∗∗∗p < 0.001. Statistical tests included unpaired Student’s t test (D,E).

Ghrelin stimulates AMPK activity in the hypothalamus (Andersson et al., 2004). However, ghrelin-induced AMPK activity was equal in primary hypothalamic neurons of *CaMK1D*^-/-^ and *CaMK1D^+/+^* mice as shown by assessment of activatory phosphorylation of AMPK and phosphorylation of its target acetyl-CoA carboxylase (ACC) (Figure 6C). Ghrelin-induced cAMP response element (CRE)-binding protein (CREB) phosphorylation promotes expression of AgRP and NPY that mediate the orexigenic action of ghrelin (Sakkou et al., 2007). Interestingly, despite higher basal expression of total CREB, levels of phosphorylated CREB were lower in control treated as well as ghrelin-treated *CaMK1D*^-/-^ cells as compared to *CaMK1D^+/+^* cells (Figure 6C). Even though activatory phosphorylation of CREB was induced upon ghrelin stimulation in *CaMK1D*^-/-^ cells, ghrelin-induced transcription of AgRP and NPY but not POMC was almost abolished in *CaMK1D*^-/-^ cells (Figure 6D), indicating that the overall reduction of activatory CREB phosphorylation constitutes at least one plausible explanation for reduced CREB-dependent expression of AgRP and NPY. In addition, AgRP immunochemistry revealed reduced levels of this neuropeptide in synaptic projections of AgRP neurons located in the PVN of *CaMK1D*^-/-^ mice under stimulatory conditions as compared to control animals (Figure 6E and F). Reduced levels and thus decreased inhibitory action of AgRP and NPY on predominantly anorexigenic neurons in the PVN is thus a likely mechanism underlying reduced food intake and body weight gain despite normal AgRP neuronal activity. Hence, CaMK1D in AgRP neurons is required for CREB-dependent expression of the orexigenic neuropeptides AgRP and NPY, thereby regulating food intake and obesity.

## Discussion

A possible role of CaMK1D in obesity and T2D has been predicted based on recent GWAS studies. However, the function of CaMK1D in physiology and metabolic disease *in vivo* was unknown thus far. In our study, using a loss-of-function approach in mice, we discovered an unpredicted role of CaMK1D in central control of food intake. We also excluded a cell-autonomous role of CaMK1D in the liver and pancreas to maintain energy homeostasis.

We found that CaMK1D is specifically required in AgRP neurons to promote ghrelin-induced hyperphagia and body weight gain. As genetic studies predicted enhanced expression of CaMK1D to contribute to T2D (Thurner et al., 2018; Xue et al., 2018), our data also fit with a model in which enhanced CaMK1D signaling in AgRP neurons promotes obesity. Deletion of *CaMK1D* in AgRP neurons is sufficient to trigger significant effects on body weight and food intake seen in global *CaMK1D* knockout mice highlighting the importance of CaMK1D signaling in this subpopulation of neurons. In line with this, CaMK1D is activated by ghrelin in NPY/AgRP neurons. Yet, we cannot fully exclude other central functions of CaMK1D signaling that may also contribute to altered body weight gain. Given the fact that AgRP neuron activity has been linked to control of liver glucose production and insulin sensitivity (Könner et al., 2007) as well as nutrient partitioning through dynamic change of the autonomic nervous system (Denis et al., 2014), it is still possible that lack of CaMK1D specifically in AgRP neuron alters insulin and glucose metabolism in addition of the feeding phenoptype. Moreover, CaMK1D is widely expressed in the CNS and ghrelin has been reported to act in different hypothalamic and extrahypothalamic areas to induce feeding. Having that said, we showed that CaMK1D is redundant in anorexigenic POMC neurons. Even though deletion of *CaMK1D* in AgRP neurons largely recapitulates phenotypes seen in whole-body knockout mice, this does not fully exclude functions in other organs implicated in energy homeostasis that we did not yet explore.

Importantly, CaMK1D is dispensable for ghrelin-stimulated electrical activity of AgRP neurons. This finding is in line with a model in which ghrelin-driven neuronal activity induces membrane depolarization and calcium changes which in turn may trigger CaMK1D activation and CaMK1D-dependent responses including CREB-dependent transcription (Sheng et al., 1991). Thus, our study has identified a so far unknown signaling pathway in AgRP neurons that links neuronal activity to CREB-dependent transcription (Figure 6G).

CREB-dependent transcription has been shown to regulate fundamental processes in neuronal development, activity-dependent dendritic outgrowth, and synaptic plasticity (Flavell and Greenberg, 2008). In AgRP neurons, CREB controls transcription of AgRP and NPY (Sakkou et al., 2007). In accordance with this finding, we found that ghrelin failed to induce AgRP and NPY transcription in *CaMK1D*-deficient hypothalamus and that the number of AgRP projections to the PVN were reduced. In fact, it has been demonstrated that ghrelin failed to stimulate feeding upon chemical and genetic inhibition of AgRP and NPY (Andrews et al., 2008; Aponte et al., 2011; Chen et al., 2004; Luquet et al., 2005; Nakazato et al., 2001). Even though AgRP is one of the crucial neuropeptides inhibiting anorexigenic neurons in the PVN, other CREB-dependent functions might be affected in *CaMK1D*-deficient AgRP neurons that may also contribute to the observed metabolic phenotypes to be investigated in the future. Moreover, CaMK1D might also regulate CREB-independent functions that need to be identified.

Central ghrelin administration induced AMPK phosphorylation and activation (Kola et al., 2005; López et al., 2008) and ghrelin responses could be alleviated through AMPK inhibition (Anderson et al., 2008). AMPK activation was dependent on calcium changes and on CaMKK 2 activation (Anderson et al., 2008). CaMKK is also known to activate CaMK1 including CaMK1D (Figure 6G). Interestingly, AMPK activation was shown to occur in the ventro-medial nucleus of the hypothalamus (VMH), since adenoviral delivery of a dominant negative isoform of AMPK into the VMH was sufficient to block ghrelin-induced food intake (Anderson et al., 2008; García et al., 2001; López et al., 2008). Therefore, it has been suggested that AgRP/NPY levels in neurons in the ARC are regulated at a presynaptic level by AMPK signaling in neurons of the VMH. We found here that lack of CaMK1D almost entirely abolished the increase in AgRP/NPY transcription in response to ghrelin. Yet, absence of CaMK1D does not affect AMPK signaling in response to ghrelin in the hypothalamus. Given that AgRP neuron-specific deletion of CaMK1D is sufficient to reduce food intake in response to ghrelin, we propose that the transcriptional control of AgRP/NPY expression primarily depends on CaMK1D signaling in AgRP neurons. Whether CaMK1D signaling occurs downstream of AMPK-dependent presynaptic mechanisms remains to be explored. Another possibility is that CaMK1D in AgRP neurons is required to integrate the trophic action of ghrelin. Indeed, neural projections arising from AgRP neurons are fully established during a critical window both during development and in the weeks around birth (Bouret et al., 2004). Trophic action of ghrelin in that window of development has been show to control ARC AgRP neurons PVN fibers growth and connection (Steculorum et al., 2015). Hence, one can hypothesize that impaired transcriptional regulation of NPY and AgRP in AgRP neurons as a consequence of CaMK1D deletion also led to developmental defects affecting post-natal hypothalamic wiring and leading to altered metabolic control.

Elevated levels of cAMP led to CREB phosphorylation at serine 133 and mutation of this site abrogated CREB-dependent reporter gene activation (Gonzalez and Montminy, 1989). Protein kinase A (PKA) is a main mediator of cAMP-dependent phosphorylation of CREB. Indeed, ghrelin was shown to increase calcium through the cAMP-PKA pathway in NPY-expressing cells in the ARC of rats (Kohno et al., 2003). However, we observe that phosphorylation of CREB depends, at least partially, on CaMK1D activity. In fact, CREB was shown to be phosphorylated *in vitro* by both kinases, PKA and CaMK1 (Sheng et al., 1991). Given that phosphorylation is reduced but not abolished in the absence of CaMK1D both kinases might be necessary to exert robust CREB phosphorylation in response to ghrelin.

mTOR-S6K1 signaling has also been demonstrated to be involved in hypothalamic regulation of food intake in response to ghrelin through regulation of CREB phosphorylation and AgRP/NPY expression (Martins et al., 2012; Stevanovic et al., 2013). However, it is unclear so far how and in which neurons mTOR-S6K1 regulates ghrelin responses. In fact, mTORC1 signaling in AgRP neurons was shown to control circadian expression of AgRP and NPY but was redundant for regulation of food intake (Albert et al., 2015).

Altogether, we uncovered a signaling mechanism that acts in AgRP neurons to control levels of AgRP and NPY, two main orexigenic neuropeptides centrally involved in promoting food intake. Uncontrolled CaMK1D signaling in AgRP neurons represents thus a valuable mechanism promoting obesity and T2D.

**Supp Figure 1. Global deletion of *CaMK1D* in mice does not affect body size and *CaMK1D* knockout mice are born without any gross abnormalities.** (A) Knockout-first targeting strategy within the *CaMK1D* loccus and generation of floxed and total knockout mice. For further details see material and methods. (B) Gender distribution of mice and (C) Mendelian ratios of born mice with indicated genetoypes. (D) Body length and (E) Tibia Length measurements of 7-week-old mice with indicated genotypes (n=6 to 7 / group).

**Supp Figure 2. CaMK1D signaling is redundant in pancreatic beta cell and liver function.** (A) Blood insulin levels during IPGTTs in mice with indicated genotypes on a HFD. IPGTT was performed after an overnight food withdrawal. (n=11 to 11 / group). (B) Glucose-stimulated insulin secretion (GSIS) in pancreatic islets isolated from mice with indicated genotypes. (n=6 to 7 / group). (C) Expression of CaMK1D protein in pancreas and brain from mice with indicated genotypes. Tissues were analyzed for CaMK1D expression by western blot. (D) Body weight gain of mice with indicated genotypes on a Chow diet (CD) or on a High Fat Diet (HFD). (n=9 to 11/ group). (E-F) Blood glucose levels during IPGTTs in mice with indicated genotypes on a CD (E) or on a HFD (F). IPGTT was performed after overnight food withdrawal. (n=9 to 11 / group). (G) Expression of CaMK1D mRNA in liver from mice with indicated genotypes by qPCR. (H) Body weight gain of mice with indicated genotypes on a CD or on a HFD. (n=9 to 11/ group). Blood glucose levels during an IPGTT in mice with indicated genotypes on a CD (I) or on a HFD (J). IPGTT was performed after overnight food withdrawal. (n=9 to 11 / group). Blood glucose levels during an ITT in mice with indicated genotypes on a CD (K) or on a HFD (L). IPGTT was performed 4H after food withdrawal. (n=6 to 10 / group).

**Supp Figure 3. Deletion of *CaMK1D* does not result in anxiety-like behavior and stress-induced anhedonia.** (A) Distance traveled each 5 min, and (B) in total. (C) Rear number each 5 min, and (D) in total. (E) Number of entries in the center or periphery of the arena. (F) Time spent in the center or periphery of the arena. (G) Average speed during an open field test. (n=9 / group). (H) Amount of sucrose and water consumed over 3 day in while sucrose and water were both available in the same time.

**Supp Figure 4. Deletion of *CaMK1D* in the Nervous Tissue reproduces phenotypes in whole-body knockout mice.** (A) Expression of CaMK1D protein in different tissues in mice. Tissues from mice with indicated genotypes were analyzed for CaMK1D expression by western blot. (B) Body weight gain of mice with indicated genotypes on a Chow diet (CD) or on a High Fat Diet (HFD). (n=10 to 15/ group). Cumulative food intake of mice with indicated genotypes on (C) CD or on (D) HFD. (n=10 to 15 / group) (*p ≤ 0.05). (E) Cumulative food intake of mice with indicated genotypes and diets. Food intake was determined 24H after fasting. (n=12 to 17 / group). (F) Cumulative food intake after ghrelin injection (30 µg / day) in mice with indicated genotypes on a CD. (n=6 to 7 / group). ∗p < 0.05, ∗∗p< 0.01 and ∗∗∗p < 0.001. Statistical tests included two-way ANOVA (B,C,D), and unpaired Student’s t test (E,F).

**Supp Figure 5. Deletion of *CaMK1D* in POMC neurons does not affect energy metabolism in mice.** (A) Verification of recombination in the CaMK1D locus in hypothalamus (Hyp), cortex (Crtx), liver, tail and white blood cells (WBC) from mice with indicated genotypes. DNA from tissues were analyzed by PCR with primers amplifying either recombined or floxed alleles, respectively. (B) Body weight gain of mice with indicated genotypes on a Chow diet (CD) or on a High Fat Diet (HFD). (n=10 to 15/ group).(C) Cumulative food intake of mice with indicated genotypes on a CD. (n=10 to 15 / group). (D) Cumulative food intake of mice with indicated genotypes on a HFD. (n=10 to 15 / group). (E) Cumulative food intake of mice with indicated genotypes and diets. Food intake was determined 24H after fasting. (n=12 to 17 / group). (F) Cumulative food intake after ghrelin injection in mice with indicated genotypes and diets. (n=6 to 7 / group). ∗p < 0.05. Statistical tests included unpaired Student’s t test (F).

## Materials and Methods

### Animals Care

Animal care and all experimental procedures done in this study were approved by the local ethical committee (Com’Eth) in compliance with the French and European legislation on care and use of laboratory animals (APAFIS#18638-2019012510177514v4). Mice were individually housed under controlled temperature at 22°C on a 12H light/dark cycle with unrestricted access to water and prescribed diet. Food was only withdrawn if required for an experiment. Body weight and food intake were determined weekly. Animals were fed with regular chow diet (CD) or high fat diet (HFD). CD contains 73.6% calories from carbohydrates, 18.1% calories from protein, and 8.4% calories from fat (SAFE® D04 from Safe) and HFD contains 20% calories from carbohydrates, 20% calories from protein, and 60% calories from fat (D12492i from Research diet®). For all experiments only male mice were used. All experiments were performed in adult mice at the age between 5-25 weeks.

### Generation of Transgenic Mice

*CaMK1D* conditional knockout and global knockout mice were generated according to the “knockout first” strategy by the Institut Clinique de la Souris (ICS, Illkirch-Graffenstaden, France). 5’ of exon 4 of the *CaMK1D* gene a SA-ßGeo-pA trapping cassette was inserted flanked by two FRT sites, which disrupts gene function (“knockout first” allele and L3 mice). Furthermore, two LoxP sites were inserted 5’ and 3’ of exon 4. The FRT-recombinase (Flp) converted the “knockout first” allele to a conditional allele (*CaMK1D^flox/flox^*), restoring gene activity (Fig S1A). The sequences of the primers used to genotype the mice and to verify Cre-mediated recombination are provided in Table S1. *CaMK1D^flox/flox^* mice were mated with Rosa26-Cre mice expressing Cre recombinase under control of the Rosa26 promoter (for global knockout) (Soriano, 1999) resulting in the deletion of the floxed exon. The breeding colonies were maintained by mating hemizygote *CaMK1D^+/-^* females to hemizygote *CaMK1D^+/-^* males. Mice were on a C57BL/6 N/J mixed background. Tissue-specific deletion of *CaMK1D* was obtained by mating floxed mice with transgenic mice expressing Cre recombinase under the control of a tissue-specific promoter and breeding colonies were maintained by mating tissue-specific promoterCre/+;*CaMK1D^flox/+^* to *CaMK1D^flox/+^* mice. All Mice were on a C57BL/6 N/J mixed background. All cre deleter mouse lines are listed in the STAR methods section.

### Blood collection and biochemical measurements

Blood samples obtained from the tail and collected in heparinized capillaries were used to measure fasted blood glucose and insulin levels. Animals were fasted at 8 am, and the samples were collected 4 hours later. At the end of the experiment, blood was collected from the retro orbital sinus, put into tubes containing 0.2 μM EDTA and 4 mM Pefabloc® SC, and centrifuged for 15 minutes at 3,000 g to separate the plasma. Plasma was stored at –80°C. Acylated Ghrelin Leptin and Insulin were measured by ELISA.

### Glucose and Insulin Tolerance Assays

GTT: After a 16H fast, animals were injected i.p. with 2 g/kg (animals on CD) or 1 g/kg (animals on HFD) dextrose in 0.9% NaCl. Blood glucose was measured prior to and 15, 30, 45, 60, 90, and 120 minutes after injections. Blood glucose values were determined in a drop of blood sampled from the tail using an automatic glucose monitor (Accu-Check; Roche Applied Science). Plasma samples were collected at 0, 15, 30 minutes for insulin measurements.

ITT: After a 5-hour fast, animals were injected i.p. with 0.75 IU/kg recombinant human insulin (Umuline; Lilly®). Blood glucose levels were measured before and 15, 30, 40, 60 and 90 minutes after injections. The glucose disappearance rate for the ITT (kITT) (percentage/minute) was calculated using the formula as previously described (Lundbaek, 1962). kITT = 0.693 × 100 / t1/2, where t1/2 was calculated from the slope of the plasma glucose concentration, considering an exponential decrement of glucose concentration during the 30 minutes after insulin administration.

### Automated Cages Phenotyping for indirect calorimetric measurements

Twenty-five weeks old mice were acclimated in metabolic chambers (TSE LabMaster System - Metabolic Phenotyping Facility, ICS) for 1 day before the start of the recordings. Mice were continuously recorded for 1 or 2 days with measurements of locomotor activity (in the xy- and z-axes), and gas exchange (O2 and CO2) every 30 min. Energy expenditure was calculated according to the manufacturer’s guidelines (PhenoMaster Software, TSE System). The respiratory quotient was estimated by calculating the ratio of CO2 production to O2 consumption. Values were corrected by metabolic mass (lean mass + 0.2 fat mass) as previously described (Even and Nadkarni, 2012). ANCOVA analysis was done as previously described (Müller et al., 2021).

### Animal length and body composition

Animal length was assessed with X-Ray MX-20 Specimen (Faxitron - Metabolic Phenotyping Facility, ICS). Digital X-ray pictures allowed the measurement of whole body and tibia size of mice. Body composition was evaluated by Quantitative Nuclear Magnetic Resonance (qNMR) using Minispec^+^ analyzer (Bruker BioSpin S.A.S., Metabolic Phenotyping Facility, ICS).

### Leptin/ghrelin responsiveness

To assess leptin sensitivity, mice received an i.p. injection of either PBS or mouse recombinant leptin (3 mg/kg) 24H after food withdrawal, their food intake were monitored 4 and 6H following the injections. The food intake after PBS injection was compared with the food intake after leptin administration. The orexigenic response to ghrelin was determined in mice that received an i.p injection of either PBS or ghrelin (1 mg/kg). Food intake was assessed 4H and 6H after injections.

### Culture of primary cells of hypothalamus

Hypothalamus were dissected from E15.5 embryos and stored on ice in Neurobasal medium (GIBCO). Tissues were incubated for 20 min in a 37°C water bath in 100U/ml papaïn (Worthinghton) and 10mg/ml DNAse I (Worthinghton). Digestion was stopped with Ovomucoïde (Worthinghton). Tissues were transferred into 1 ml of adult neuronal growth medium consisting of Neurobasal-A medium, 3mM L-glutamine (Gibco), 1x B-27 supplement, 1x N2 supplement and antibiotics. Tissues were gently triturated until uniform cellular dissociation was achieved. Cells were counted and plated into cell culture plates coated with poly-L-lysine (Gibco).

### Western blotting

Cells were washed with ice-cold PBS on ice and snap-frozen in liquid nitrogen. Cell lysates for WB were prepared using 1x Laemmli buffer (50 mM Tris-HCl pH6.8, 100 mM DTT, 8% SDS, 0,01% bromophenol blue, 10% glycerol) supplemented with phosphatase/protease inhibitors (Cell Signaling Technology) and incubated on ice for 10 minutes. After centrifugation at 16 000 g for 10 minutes at 4°C cleared supernatant was transferred to the new tubes and was used immediately stored at −80°C until used. Total protein was measured using the BCA method by Pierce™ BCA Protein Assay Kit (ThermoFisher). Samples (20-50 μg of total protein content) were boiled and resolved on 10% acrylamide gels using standard Tris-Glycine SDS-PAGE or Phostag gels. Proteins were transferred to PVDF membranes (Millipore) and blotted with antibodies listed in the Antibodies section. For membrane blocking and primary antibody dilution 5% BSA (w/v) in TBST was used. All incubations with primary antibodies were performed for 16 hours at 4°C. Blots were developed using SuperSignal West Pico (Pierce, Ref. 34580) or Luminata Forte Western HRP substrate (Merck Millipore, Ref. WBLUF0500).

### Hypothalamic mRNA quantification

Total RNA from hypothalamus was extracted using an RNeasy Lipid Tissue Mini Kit (QIAGEN) and quantified spectrophotometrically. Single-stranded cDNA was synthesized using SuperScript IV RNase Reverse Transcriptase (Invitrogen) according to the manufacturer’s directions. Real-time PCR was carried out using an LightCycler® 480 (Roche) with Fast SYBR® Green Master Mix (Roche) and the primers listed in the primers section. Quantifications were done according the Pfaffl method (Pfaffl, 2001).

### Immunohistochemistry

Mediobasal hypothalamic sections from brains were prefixed with paraformaldehyde during 24h and incubated in 30% sucrose (Fisher Scientific) 24H at 4°C. Brains were embedded in OCT, frozen at −80°C and stored at −80°C. 30 µm-thick sections were cut with a cryostat (Leica CM3050 S, France), stored at 4°C in sodium phosphate buffer. Sections were processed as follows: Day 1: free-floating sections were rinsed in PBS, incubated for 20 min in PBS containing 0.3% Tween-20, and then rinsed three times for 10 min each in PBS. Slices were incubated 1h with 5% donkey serum in 0.3% PBS-T and then overnight or 72H at 4°C with the primary antibodies described in the antibodies section. Slides were rinsed three times for 10 min in 0.3% PBS-T and incubated for 60 min with secondary antibodies. Sections were rinsed three times for 10 min in PBS before mounting. Tissues were observed on a confocal laser scanning microscope, TCS SP8X; with Leica software LAS X navigator, using a HC PL APO CS2 20x /0.75 dry leica objective. The objectives and the pinhole setting (1 airy unit, au) remained unchanged during the acquisition of a series for all images. Quantification of immuno-positive cells was performed using the cell counter plugin of the ImageJ software taking as standard reference a fixed threshold of fluorescence.

### Electrophysiology

Experiments were performed on brain slices from 9-12 week old male CaMK1D+/+ and CaMK1D-/- mice that expressed enhanced green fluorescent protein (EGFP) selectively in AgRP neurons. Animals were kept under standard laboratory conditions, with tap water and chow available ad libitum, on a 12h light/dark cycle. The animals were lightly anesthetized with isoflurane (B506; AbbVie Deutschland GmbH and Co KG, Ludwigshafen, Germany) and decapitated. Coronal slices (280 µm) containing the arcuate nucleus of the hypothalamus were cut with a vibration microtome (VT1200 S; Leica, Germany) under cold (4°C), carbogenated (95% O2 and 5% CO2), glycerol-based modified artificial cerebrospinal fluid (GaCSF) (Ye et al., 2006).

Current-clamp recordings of GFP-expressing AgRP neurons were performed at ∼32°C. Neurons were visualized with a fixed stage upright microscope (Zeiss AxioExaminer, Zeiss, Germany) using 40x water-immersion objective (W Plan-Apochromat 40x/1.0 DIC M27, 1 numerical aperture, 2.5 mm working distance; Zeiss) with infrared differential interference contrast optics (Dodt and Zieglgänsberger, 1990) and fluorescence optics. GFP-expressing AgRP neurons were identified by their anatomical location in the arcuate nucleus and by their fluorescent label. Perforated patch experiments were conducted using protocols modified from (Horn and Marty, 1988) and (Akaike and Harata, 1994).

The spontaneous firing frequency was measured for 5 min after perforation. To measure intrinsic electrophysiological properties series of hyperpolarizing and depolarizing current pulses were applied under current clamp from a membrane potential of ∼-70 mV. For input resistance and capacitance measurements, hyperpolarizing current steps with −2 pA increments were applied. For excitability measurements, depolarizing 1 s current steps with +2 pA increments were applied. The specific protocols are given in Results.

To investigate the modulatory effect of ghrelin (031-31, Phoenix Pharmaceuticals), 100 nM ghrelin was bath applied for 8-10 min. The ghrelin effect was analyzed by comparing the action potential frequencies that were measured during 2 min intervals that were recorded directly before and at the end of the peptide applications.

### Data analysis of electrophysiological data

Data analysis was performed with Spike2 (version 7; Cambridge Electronic Design Ltd., Cambridge, UK), Igor Pro 6 (Wavemetrics, Portland, OR, USA), and Graphpad Prism 8. If not stated otherwise, all calculated values are expressed as means ± SEM (standard error of the mean). The horizontal lines show the data’s median. The whiskers were calculated according to the ‘Tukey’ method. For comparisons of independent nonparametric distributions, the Mann-Whitney-U-test was used. Linear regressions were compared using the F-test. A significance level of 0.05 was accepted for all tests. Exact p-values are reported if p > 0.05. In the figures, n values are given in brackets.

### Quantification and statistical analysis

All statistical comparisons were performed with Prism 6 (GraphPad Software, La Jolla, CA, USA) or R software for ANCOVA analysis. The statistical tests used are listed along with the statistical values in the Supplemental Tables. All the data were analyzed using either Student t test (paired or unpaired) with equal variances or One-way ANOVA or Two-way ANOVA. In all cases, significance threshold was automatically set at p < 0.05. ANOVA analyses were followed by Bonferroni post hoc test for specific comparisons only when overall ANOVA revealed a significant difference (at least p < 0.05).

## STAR METHODS

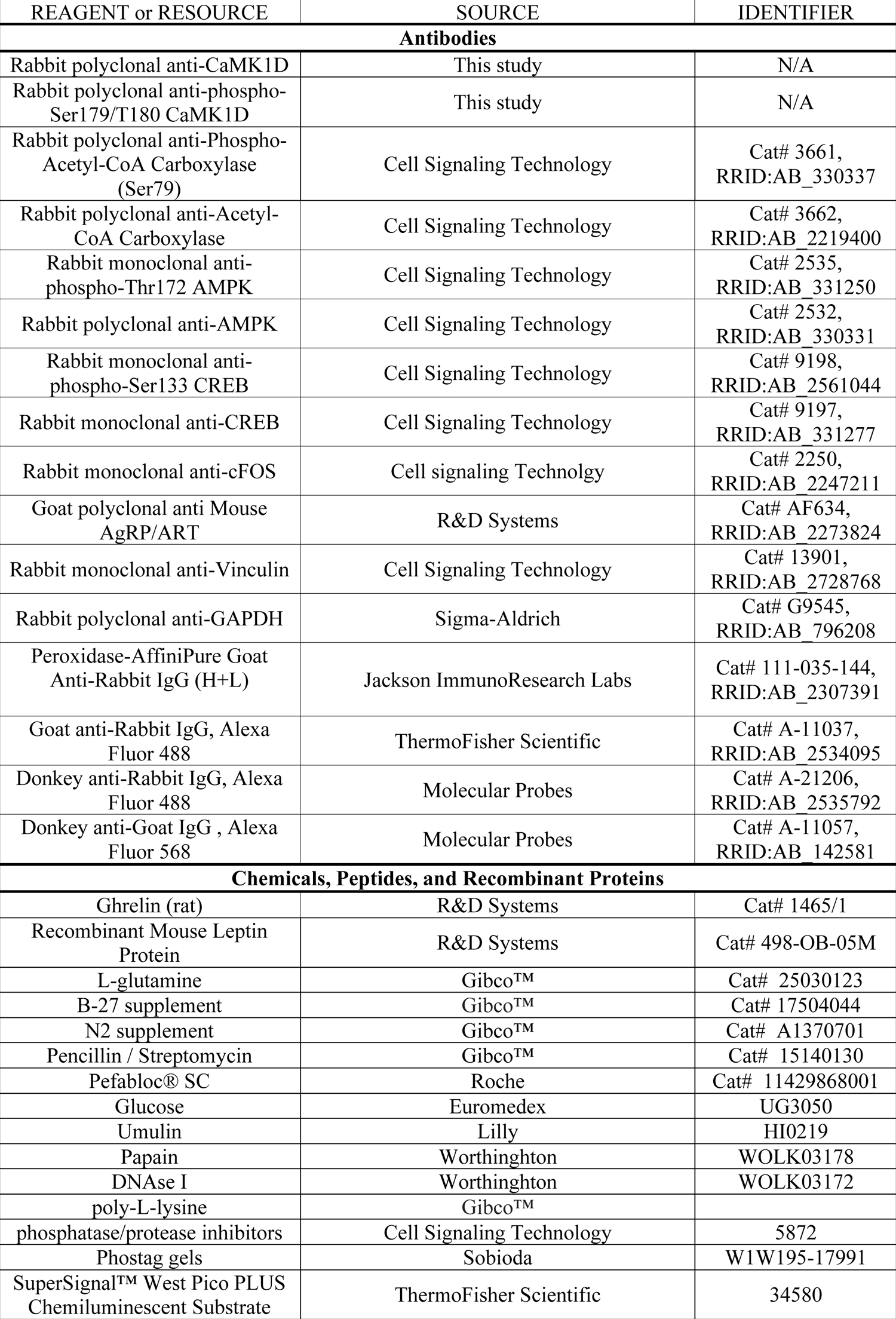
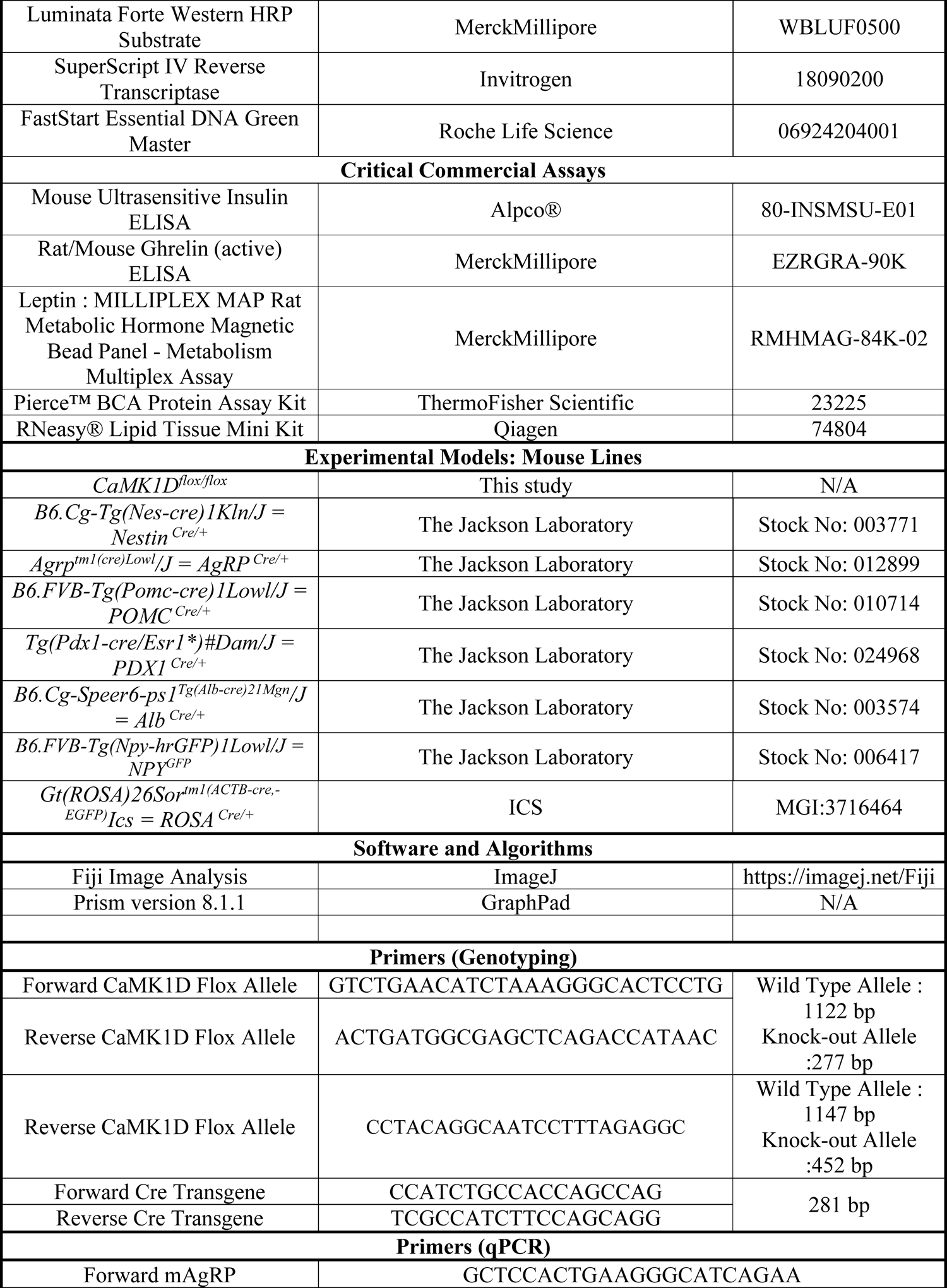
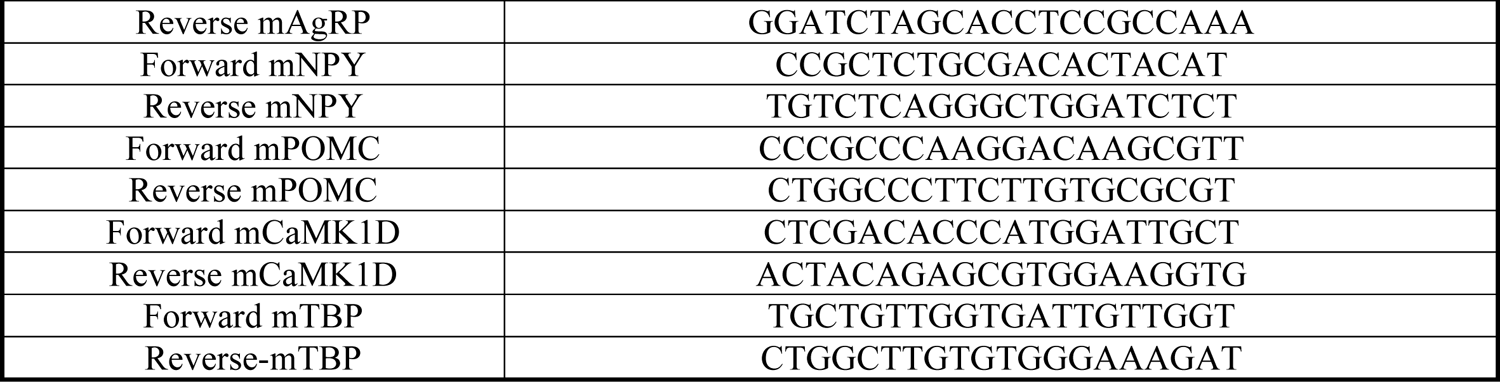

## Supporting information

Vivot et al._2021_Supp Figures

## Acknowledgments

We thank T. Alquier at University of Montreal, E. Pangou and D. Dembele at the IGBMC for helpful discussions. We thank the Imaging Center of the IGBMC (ICI) and the IGBMC core facilities for their support on this research. We are grateful to F. Berditchevski at Nottingham University for the pS179/Thr180 CaMK1D antibody. This work was supported by the Agence Nationale de la Recherche (ANR) (AAPG 2017 LYSODIABETES and AAPG 2021 HypoCaMK)), by the Fondation de Recherche Médicale (FRM) – Program: Equipe FRM (EQU201903007859, Prix Roger PROPICE pour la recherche sur le cancer du pancréas), by the FHU-OMAGE of region Grand-Est, from the European Foundation for the Study of Diabetes (EFSD)/Novo Nordisk Diabetes Research Programme and by the ANR-10-LABX-0030-INRT grant as well as the ANR-11-INBS-0009-INGESTEM grant, both French State funds managed by the ANR under the frame program Investissements d’Avenir. K. Vivot was supported by an Individual Fellowship (798961 INSULYSOSOME) in the framework of the Marie-Sklodovska Curie actions of the European Commission. G. Yeghiazaryan received financial doctoral support from DFG-233886668/RTG1960.

## Author contributions

Conceptualization: R.R., R.P.N, P.K and S.L., Software: E.G., Methodology: K.V., Z.Z, G.Y., E.E., E.C.C. and M.Q, Validation: K.V., Formal Analysis: K.V., C.M., Investigation: K.V., G.M., Z.Z., E.E., G.Y., M.Q., A.S., M.F and C.M., Resources: E.E., Writing-Original Draft: R.R. and K.V., Supervision: S.L., P.K., R.P.R., A.C., and R.R, Funding Acquisition: R.R. and S.L.

## References

1. Akaike, N., and Harata, N. (1994). Nystatin perforated patch recording and its applications to analyses of intracellular mechanisms. Jpn J Physiol 44, 433–473.

2. Albert, V., Cornu, M., and Hall, M.N. (2015). mTORC1 signaling in Agrp neurons mediates circadian expression of Agrp and NPY but is dispensable for regulation of feeding behavior. Biochem Biophys Res Commun 464, 480–486.

3. Anderson, K.A., Ribar, T.J., Lin, F., Noeldner, P.K., Green, M.F., Muehlbauer, M.J., Witters, L.A., Kemp, B.E., and Means, A.R. (2008). Hypothalamic CaMKK2 contributes to the regulation of energy balance. Cell Metab. 7, 377–388.

4. Andersson, U., Filipsson, K., Abbott, C.R., Woods, A., Smith, K., Bloom, S.R., Carling, D., and Small, C.J. (2004). AMP-activated protein kinase plays a role in the control of food intake. J Biol Chem 279, 12005–12008.

5. Andrews, Z.B., Liu, Z.-W., Walllingford, N., Erion, D.M., Borok, E., Friedman, J.M., Tschöp, M.H., Shanabrough, M., Cline, G., Shulman, G.I., et al. (2008). UCP2 mediates ghrelin’s action on NPY/AgRP neurons by lowering free radicals. Nature 454, 846–851.

6. Aponte, Y., Atasoy, D., and Sternson, S.M. (2011). AGRP neurons are sufficient to orchestrate feeding behavior rapidly and without training. Nat Neurosci 14, 351–355.

7. Bonnefond, A., and Froguel, P. (2015). Rare and common genetic events in type 2 diabetes: what should biologists know? Cell Metab. 21, 357–368.

8. Bouret, S.G., Draper, S.J., and Simerly, R.B. (2004). Formation of projection pathways from the arcuate nucleus of the hypothalamus to hypothalamic regions implicated in the neural control of feeding behavior in mice. J Neurosci 24, 2797–2805.

9. Buchser, W.J., Slepak, T.I., Gutierrez-Arenas, O., Bixby, J.L., and Lemmon, V.P. (2010). Kinase/phosphatase overexpression reveals pathways regulating hippocampal neuron morphology. Mol. Syst. Biol. 6, 391.

10. Chen, H.Y., Trumbauer, M.E., Chen, A.S., Weingarth, D.T., Adams, J.R., Frazier, E.G., Shen, Z., Marsh, D.J., Feighner, S.D., Guan, X.-M., et al. (2004). Orexigenic action of peripheral ghrelin is mediated by neuropeptide Y and agouti-related protein. Endocrinology 145, 2607–2612.

11. Chen, S.-R., Chen, H., Zhou, J.-J., Pradhan, G., Sun, Y., Pan, H.-L., and Li, D.-P. (2017). Ghrelin receptors mediate ghrelin-induced excitation of agouti-related protein/neuropeptide Y but not pro-opiomelanocortin neurons. J. Neurochem. 142, 512–520.

12. Denis, R.G.P., Joly-Amado, A., Cansell, C., Castel, J., Martinez, S., Delbes, A.S., and Luquet, S. (2014). Central orchestration of peripheral nutrient partitioning and substrate utilization: implications for the metabolic syndrome. Diabetes Metab 40, 191–197.

13. Dodt, H.U., and Zieglgänsberger, W. (1990). Visualizing unstained neurons in living brain slices by infrared DIC-videomicroscopy. Brain Res 537, 333–336.

14. Even, P.C., and Nadkarni, N.A. (2012). Indirect calorimetry in laboratory mice and rats: principles, practical considerations, interpretation and perspectives. Am J Physiol Regul Integr Comp Physiol 303, R459–476.

15. Flavell, S.W., and Greenberg, M.E. (2008). Signaling mechanisms linking neuronal activity to gene expression and plasticity of the nervous system. Annu Rev Neurosci 31, 563–590.

16. García, A., Alvarez, C.V., Smith, R.G., and Diéguez, C. (2001). Regulation of Pit-1 expression by ghrelin and GHRP-6 through the GH secretagogue receptor. Mol Endocrinol 15, 1484–1495.

17. Gonzalez, G.A., and Montminy, M.R. (1989). Cyclic AMP stimulates somatostatin gene transcription by phosphorylation of CREB at serine 133. Cell 59, 675–680.

18. Gu, G., Dubauskaite, J., and Melton, D.A. (2002). Direct evidence for the pancreatic lineage: NGN3+ cells are islet progenitors and are distinct from duct progenitors. Development 129, 2447–2457.

19. Haney, S., Zhao, J., Tiwari, S., Eng, K., Guey, L.T., and Tien, E. (2013). RNAi screening in primary human hepatocytes of genes implicated in genome-wide association studies for roles in type 2 diabetes identifies roles for CAMK1D and CDKAL1, among others, in hepatic glucose regulation. PLoS ONE 8, e64946.

20. Haribabu, B., Hook, S.S., Selbert, M.A., Goldstein, E.G., Tomhave, E.D., Edelman, A.M., Snyderman, R., and Means, A.R. (1995). Human calcium-calmodulin dependent protein kinase I: cDNA cloning, domain structure and activation by phosphorylation at threonine-177 by calcium-calmodulin dependent protein kinase I kinase. EMBO J 14, 3679–3686.

21. Hoffman, G.E., Smith, M.S., and Verbalis, J.G. (1993). c-Fos and related immediate early gene products as markers of activity in neuroendocrine systems. Front Neuroendocrinol 14, 173–213.

22. Horn, R., and Marty, A. (1988). Muscarinic activation of ionic currents measured by a new whole-cell recording method. J Gen Physiol 92, 145–159.

23. Jais, A., and Brüning, J.C. (2021). Arcuate nucleus-dependent regulation of metabolism - pathways to obesity and diabetes mellitus. Endocr Rev bnab025.

24. Kamata, A., Sakagami, H., Tokumitsu, H., Owada, Y., Fukunaga, K., and Kondo, H. (2007). Spatiotemporal expression of four isoforms of Ca2+/calmodulin-dependent protein kinase I in brain and its possible roles in hippocampal dendritic growth. Neurosci. Res. 57, 86–97.

25. Kim, K.-S., Seeley, R.J., and Sandoval, D.A. (2018). Signalling from the periphery to the brain that regulates energy homeostasis. Nat. Rev. Neurosci. 19, 185–196.

26. Kohno, D., Gao, H.-Z., Muroya, S., Kikuyama, S., and Yada, T. (2003). Ghrelin directly interacts with neuropeptide-Y-containing neurons in the rat arcuate nucleus: Ca2+ signaling via protein kinase A and N-type channel-dependent mechanisms and cross-talk with leptin and orexin. Diabetes 52, 948–956.

27. Kola, B., Hubina, E., Tucci, S.A., Kirkham, T.C., Garcia, E.A., Mitchell, S.E., Williams, L.M., Hawley, S.A., Hardie, D.G., Grossman, A.B., et al. (2005). Cannabinoids and ghrelin have both central and peripheral metabolic and cardiac effects via AMP-activated protein kinase. J Biol Chem 280, 25196–25201.

28. Könner, A.C., Janoschek, R., Plum, L., Jordan, S.D., Rother, E., Ma, X., Xu, C., Enriori, P., Hampel, B., Barsh, G.S., et al. (2007). Insulin action in AgRP-expressing neurons is required for suppression of hepatic glucose production. Cell Metab 5, 438–449.

29. Kooner, J.S., Saleheen, D., Sim, X., Sehmi, J., Zhang, W., Frossard, P., Been, L.F., Chia, K.-S., Dimas, A.S., Hassanali, N., et al. (2011). Genome-wide association study in individuals of South Asian ancestry identifies six new type 2 diabetes susceptibility loci. Nat. Genet. 43, 984–989.

30. Locke, A.E., Kahali, B., Berndt, S.I., Justice, A.E., Pers, T.H., Day, F.R., Powell, C., Vedantam, S., Buchkovich, M.L., Yang, J., et al. (2015). Genetic studies of body mass index yield new insights for obesity biology. Nature 518, 197–206.

31. López, M., Lage, R., Saha, A.K., Pérez-Tilve, D., Vázquez, M.J., Varela, L., Sangiao-Alvarellos, S., Tovar, S., Raghay, K., Rodríguez-Cuenca, S., et al. (2008). Hypothalamic fatty acid metabolism mediates the orexigenic action of ghrelin. Cell Metab 7, 389–399.

32. Lundbaek, K. (1962). Intravenous glucose tolerance as a tool in definition and diagnosis of diabetes mellitus. Br Med J 1, 1507–1513.

33. Luquet, S., Perez, F.A., Hnasko, T.S., and Palmiter, R.D. (2005). NPY/AgRP neurons are essential for feeding in adult mice but can be ablated in neonates. Science 310, 683–685.

34. Martins, L., Fernández-Mallo, D., Novelle, M.G., Vázquez, M.J., Tena-Sempere, M., Nogueiras, R., López, M., and Diéguez, C. (2012). Hypothalamic mTOR signaling mediates the orexigenic action of ghrelin. PLoS One 7, e46923.

35. Morris, A.P., Voight, B.F., Teslovich, T.M., Ferreira, T., Segrè, A.V., Steinthorsdottir, V., Strawbridge, R.J., Khan, H., Grallert, H., Mahajan, A., et al. (2012). Large-scale association analysis provides insights into the genetic architecture and pathophysiology of type 2 diabetes. Nat. Genet. 44, 981–990.

36. Morton, G.J., Cummings, D.E., Baskin, D.G., Barsh, G.S., and Schwartz, M.W. (2006). Central nervous system control of food intake and body weight. Nature 443, 289–295.

37. Müller, T.D., Nogueiras, R., Andermann, M.L., Andrews, Z.B., Anker, S.D., Argente, J., Batterham, R.L., Benoit, S.C., Bowers, C.Y., Broglio, F., et al. (2015). Ghrelin. Mol Metab 4, 437–460.

38. Müller, T.D., Klingenspor, M., and Tschöp, M.H. (2021). Revisiting energy expenditure: how to correct mouse metabolic rate for body mass. Nat Metab 3, 1134–1136.

39. Nakazato, M., Murakami, N., Date, Y., Kojima, M., Matsuo, H., Kangawa, K., and Matsukura, S. (2001). A role for ghrelin in the central regulation of feeding. Nature 409, 194– 198.

40. Pfaffl, M.W. (2001). A new mathematical model for relative quantification in real-time RT-PCR. Nucleic Acids Res 29, e45.

41. Postic, C., Shiota, M., Niswender, K.D., Jetton, T.L., Chen, Y., Moates, J.M., Shelton, K.D., Lindner, J., Cherrington, A.D., and Magnuson, M.A. (1999). Dual roles for glucokinase in glucose homeostasis as determined by liver and pancreatic beta cell-specific gene knock-outs using Cre recombinase. J Biol Chem 274, 305–315.

42. Sakkou, M., Wiedmer, P., Anlag, K., Hamm, A., Seuntjens, E., Ettwiller, L., Tschöp, M.H., and Treier, M. (2007). A role for brain-specific homeobox factor Bsx in the control of hyperphagia and locomotory behavior. Cell Metab 5, 450–463.

43. Schmitt, J.M., Guire, E.S., Saneyoshi, T., and Soderling, T.R. (2005). Calmodulin-dependent kinase kinase/calmodulin kinase I activity gates extracellular-regulated kinase-dependent long-term potentiation. J. Neurosci. 25, 1281–1290.

44. Senga, Y., Ishida, A., Shigeri, Y., Kameshita, I., and Sueyoshi, N. (2015). The Phosphatase-Resistant Isoform of CaMKI, Ca2+/Calmodulin-Dependent Protein Kinase Iδ (CaMKIδ), Remains in Its “Primed” Form without Ca2+ Stimulation. Biochemistry 54, 3617–3630.

45. Sheng, M., Thompson, M.A., and Greenberg, M.E. (1991). CREB: a Ca(2+)-regulated transcription factor phosphorylated by calmodulin-dependent kinases. Science 252, 1427– 1430.

46. Shu, X.O., Long, J., Cai, Q., Qi, L., Xiang, Y.-B., Cho, Y.S., Tai, E.S., Li, X., Lin, X., Chow, W.-H., et al. (2010). Identification of new genetic risk variants for type 2 diabetes. PLoS Genet. 6, e1001127.

47. Simonis-Bik, A.M., Nijpels, G., van Haeften, T.W., Houwing-Duistermaat, J.J., Boomsma, D.I., Reiling, E., van Hove, E.C., Diamant, M., Kramer, M.H.H., Heine, R.J., et al. (2010). Gene variants in the novel type 2 diabetes loci CDC123/CAMK1D, THADA, ADAMTS9, BCL11A, and MTNR1B affect different aspects of pancreatic beta-cell function. Diabetes 59, 293–301.

48. Soriano, P. (1999). Generalized lacZ expression with the ROSA26 Cre reporter strain. Nat Genet 21, 70–71.

49. Steculorum, S.M., Collden, G., Coupe, B., Croizier, S., Lockie, S., Andrews, Z.B., Jarosch, F., Klussmann, S., and Bouret, S.G. (2015). Neonatal ghrelin programs development of hypothalamic feeding circuits. J Clin Invest 125, 846–858.

50. Stevanovic, D., Trajkovic, V., Müller-Lühlhoff, S., Brandt, E., Abplanalp, W., Bumke-Vogt, C., Liehl, B., Wiedmer, P., Janjetovic, K., Starcevic, V., et al. (2013). Ghrelin-induced food intake and adiposity depend on central mTORC1/S6K1 signaling. Mol Cell Endocrinol 381, 280–290.

51. Stuber, G.D., and Wise, R.A. (2016). Lateral hypothalamic circuits for feeding and reward. Nat Neurosci 19, 198–205.

52. Takemoto-Kimura, S., Ageta-Ishihara, N., Nonaka, M., Adachi-Morishima, A., Mano, T., Okamura, M., Fujii, H., Fuse, T., Hoshino, M., Suzuki, S., et al. (2007). Regulation of dendritogenesis via a lipid-raft-associated Ca2+/calmodulin-dependent protein kinase CLICK-III/CaMKIgamma. Neuron 54, 755–770.

53. Thurner, M., van de Bunt, M., Torres, J.M., Mahajan, A., Nylander, V., Bennett, A.J., Gaulton, K.J., Barrett, A., Burrows, C., Bell, C.G., et al. (2018). Integration of human pancreatic islet genomic data refines regulatory mechanisms at Type 2 Diabetes susceptibility loci. Elife 7.

54. Timper, K., and Brüning, J.C. (2017). Hypothalamic circuits regulating appetite and energy homeostasis: pathways to obesity. Dis Model Mech 10, 679–689.

55. Verploegen, S., Ulfman, L., van Deutekom, H.W.M., van Aalst, C., Honing, H., Lammers, J.-W.J., Koenderman, L., and Coffer, P.J. (2005). Characterization of the role of CaMKI-like kinase (CKLiK) in human granulocyte function. Blood 106, 1076–1083.

56. Waterson, M.J., and Horvath, T.L. (2015). Neuronal Regulation of Energy Homeostasis: Beyond the Hypothalamus and Feeding. Cell Metab. 22, 962–970.

57. Wayman, G.A., Kaech, S., Grant, W.F., Davare, M., Impey, S., Tokumitsu, H., Nozaki, N., Banker, G., and Soderling, T.R. (2004). Regulation of axonal extension and growth cone motility by calmodulin-dependent protein kinase I. J. Neurosci. 24, 3786–3794.

58. Xue, A., Wu, Y., Zhu, Z., Zhang, F., Kemper, K.E., Zheng, Z., Yengo, L., Lloyd-Jones, L.R., Sidorenko, J., Wu, Y., et al. (2018). Genome-wide association analyses identify 143 risk variants and putative regulatory mechanisms for type 2 diabetes. Nat Commun 9, 2941.

59. Ye, J.H., Zhang, J., Xiao, C., and Kong, J.-Q. (2006). Patch-clamp studies in the CNS illustrate a simple new method for obtaining viable neurons in rat brain slices: glycerol replacement of NaCl protects CNS neurons. J Neurosci Methods 158, 251–259.

60. Zeggini, E., Scott, L.J., Saxena, R., Voight, B.F., Marchini, J.L., Hu, T., de Bakker, P.I., Abecasis, G.R., Almgren, P., Andersen, G., et al. (2008). Meta-analysis of genome-wide association data and large-scale replication identifies additional susceptibility loci for type 2 diabetes. Nature Genetics 40, 638–645.

