## Supplementary figures and images for "CaMK1D signaling in AgRP neurons promotes ghrelin-mediated food intake"

### Vivot et al._2021_Supp Figures

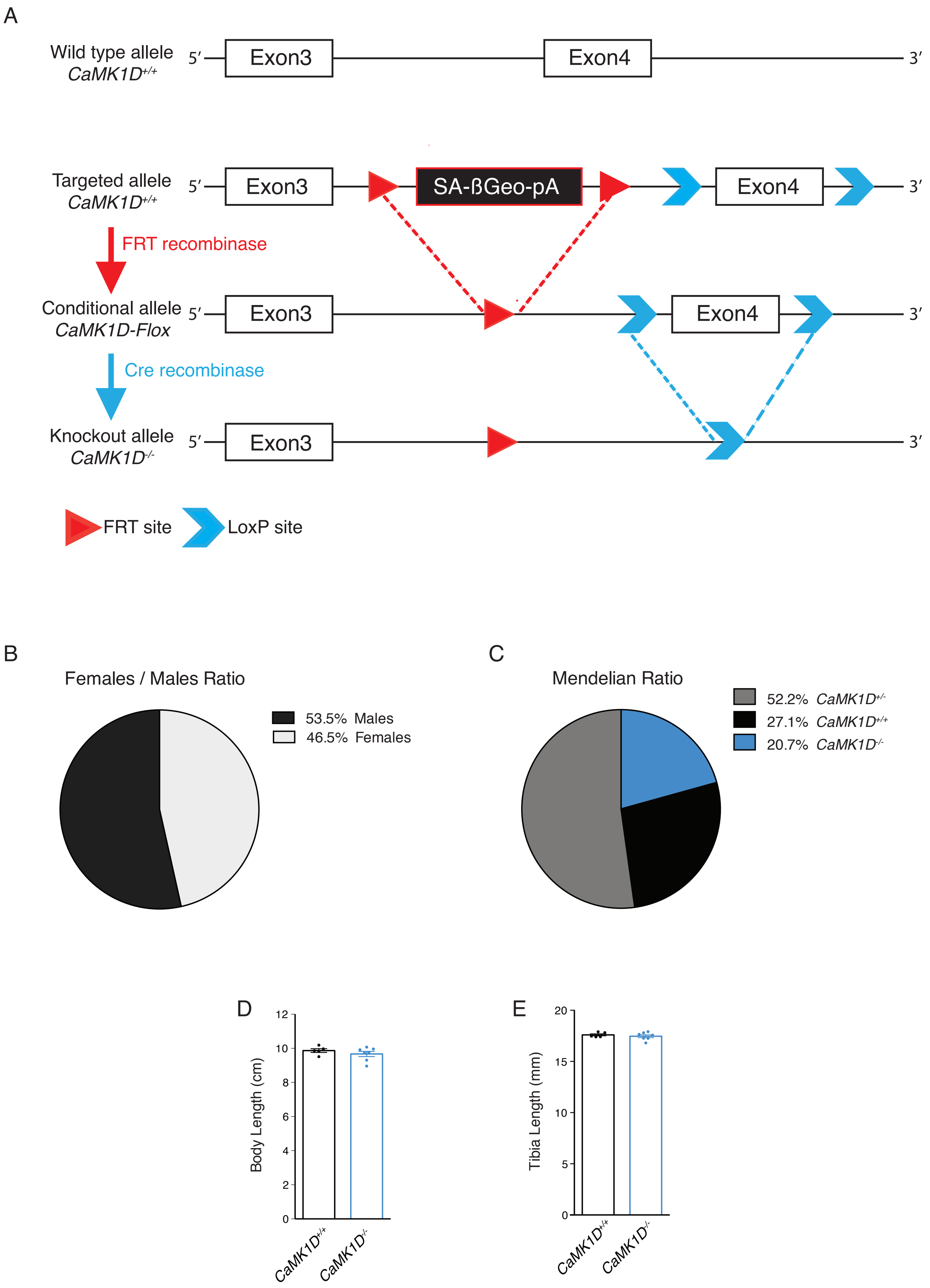

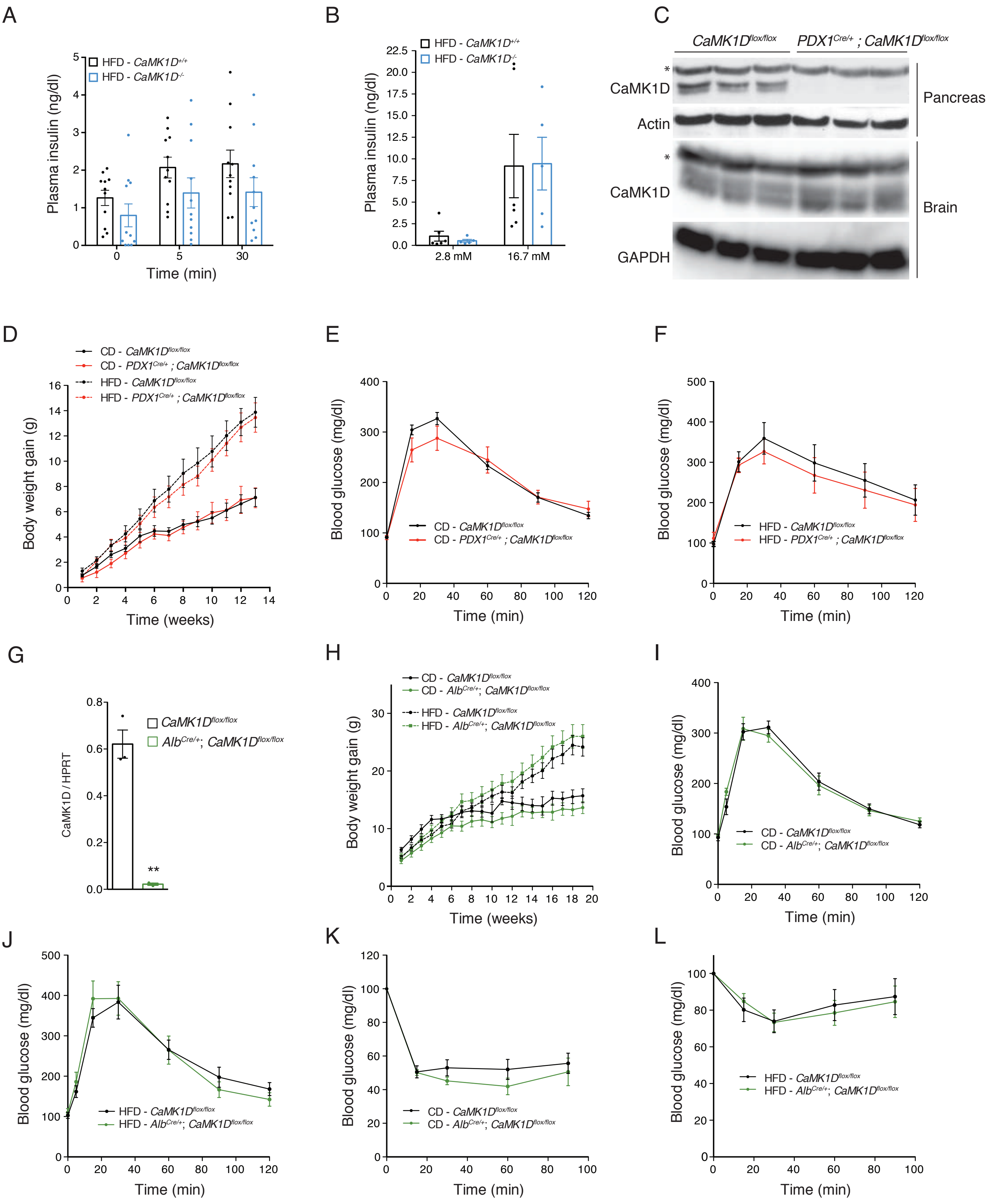

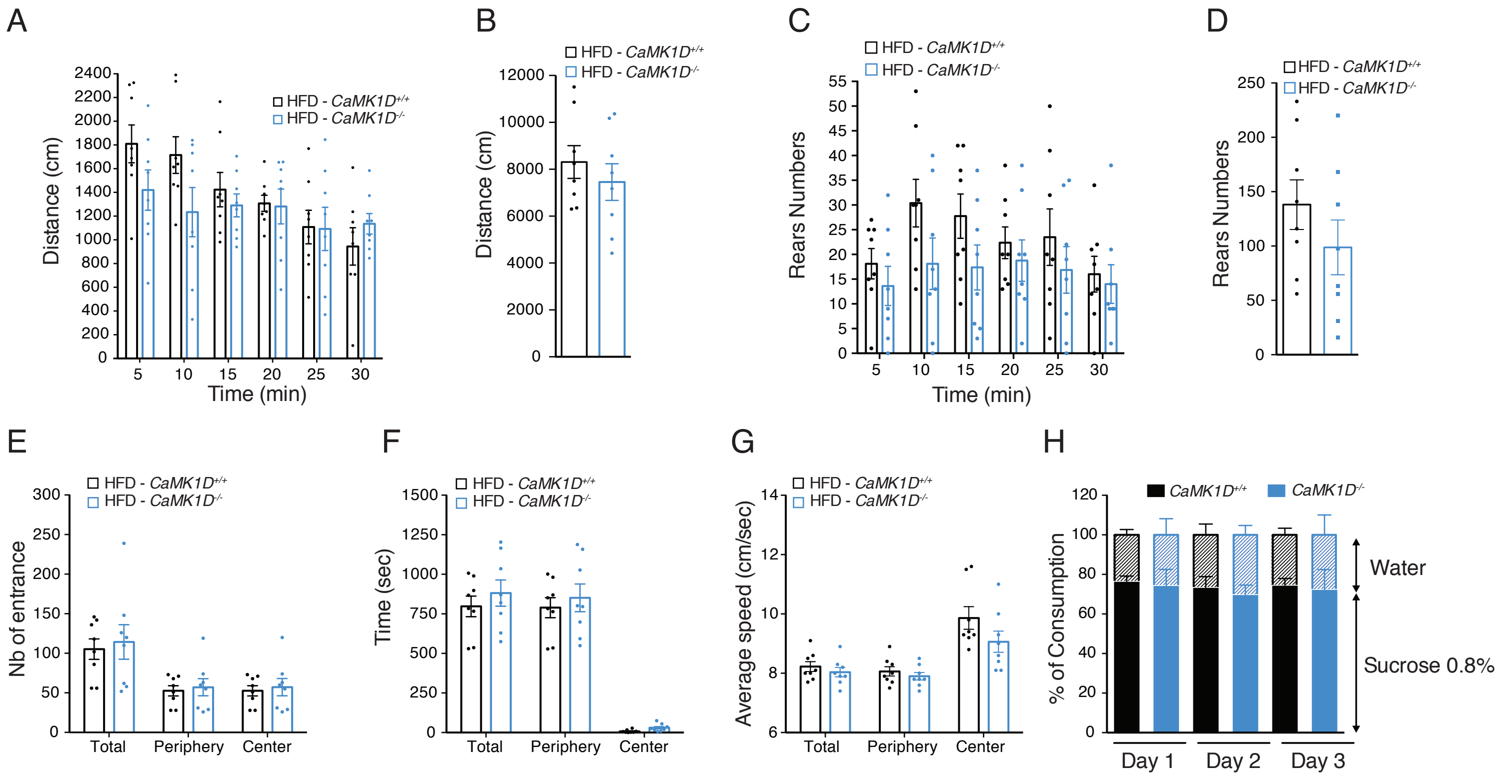

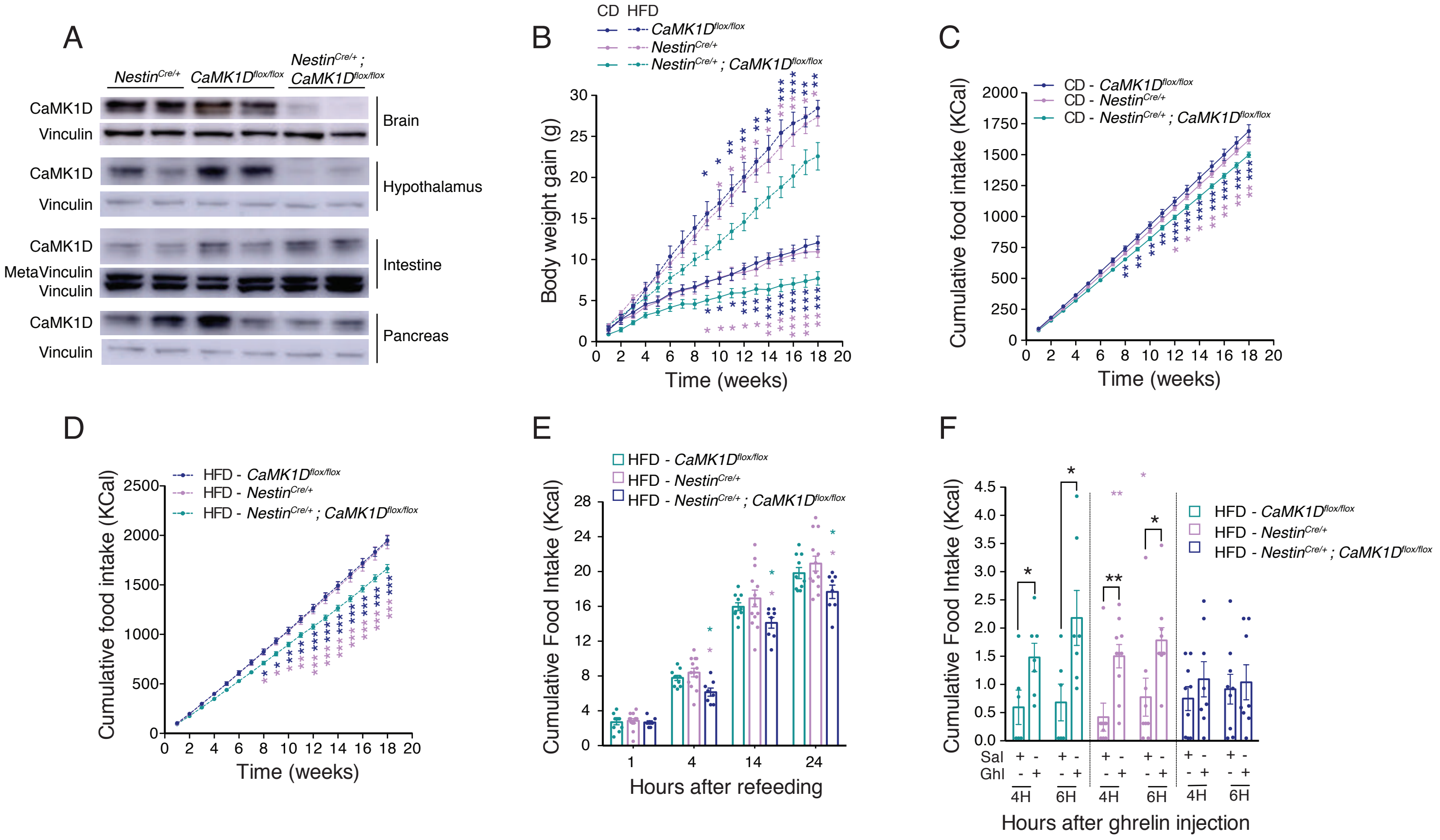

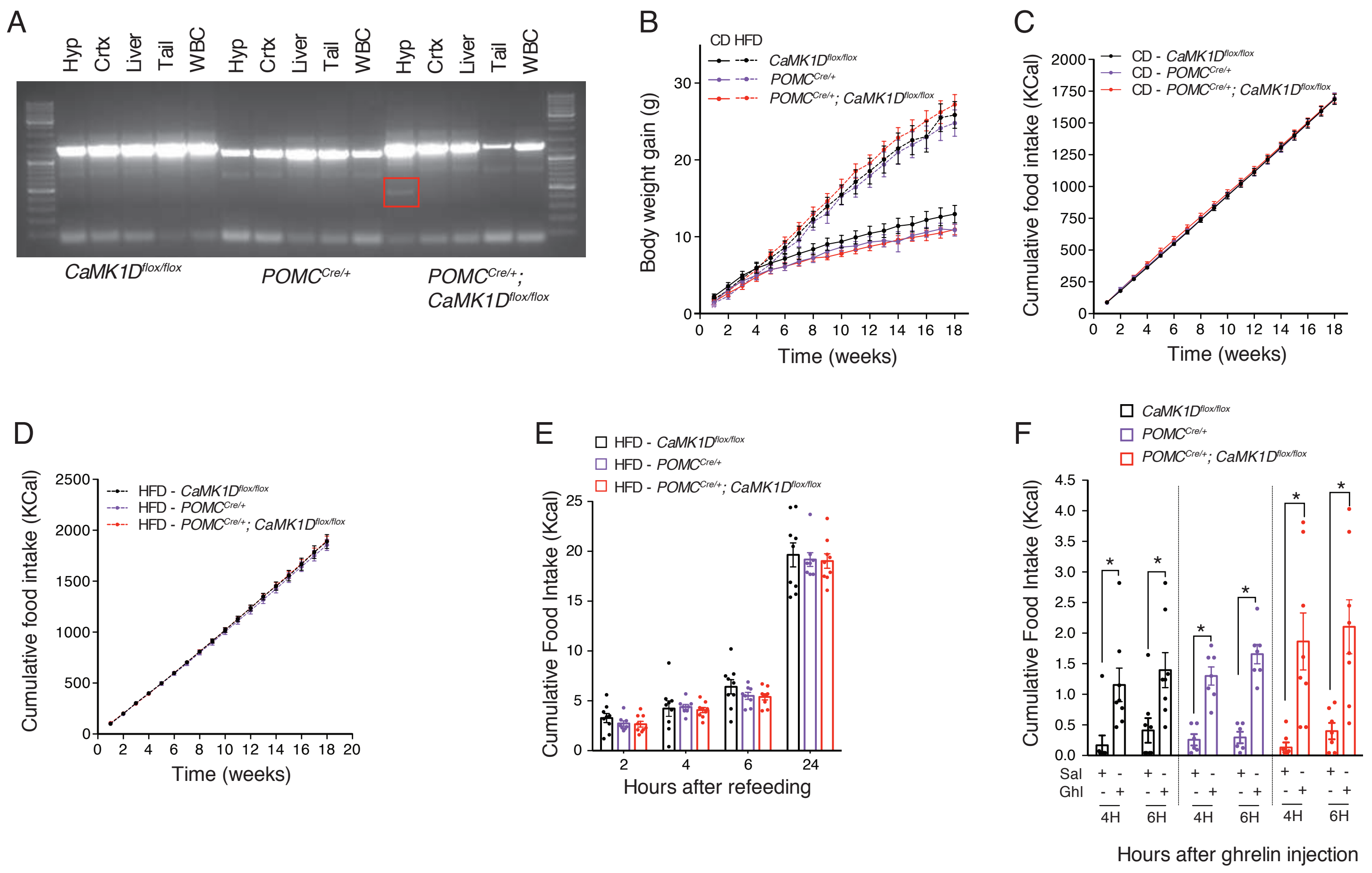
